# Antimicrobial peptides modulate lung injury by altering the intestinal microbiota

**DOI:** 10.1101/2023.03.14.529700

**Authors:** Ahmed Abdelgawad, Teodora Nicola, Isaac Martin, Brian A. Halloran, Kosuke Tanaka, Comfort Y. Adegboye, Pankaj Jain, Changchun Ren, Charitharth V. Lal, Namasivayam Ambalavanan, Amy E. O’Connell, Tamás Jilling, Kent A. Willis

## Abstract

Mammalian mucosal barriers secrete antimicrobial peptides (AMPs) as critical host-derived regulators of the microbiota. However, mechanisms that support homeostasis of the microbiota in response to inflammatory stimuli such as supraphysiologic oxygen remain unclear. Here, we show that neonatal mice breathing supraphysiologic oxygen or direct exposure of intestinal organoids to supraphysiologic oxygen suppress the intestinal expression of AMPs and alters the composition of the intestinal microbiota. Oral supplementation of the prototypical AMP lysozyme to hyperoxia exposed neonatal mice reduced hyperoxia-induced alterations in their microbiota and was associated with decreased lung injury. Our results identify a gut-lung axis driven by intestinal AMP expression and mediated by the intestinal microbiota that is linked to lung injury. Together, these data support that intestinal AMPs modulate lung injury and repair.

**Graphical Abstract:** 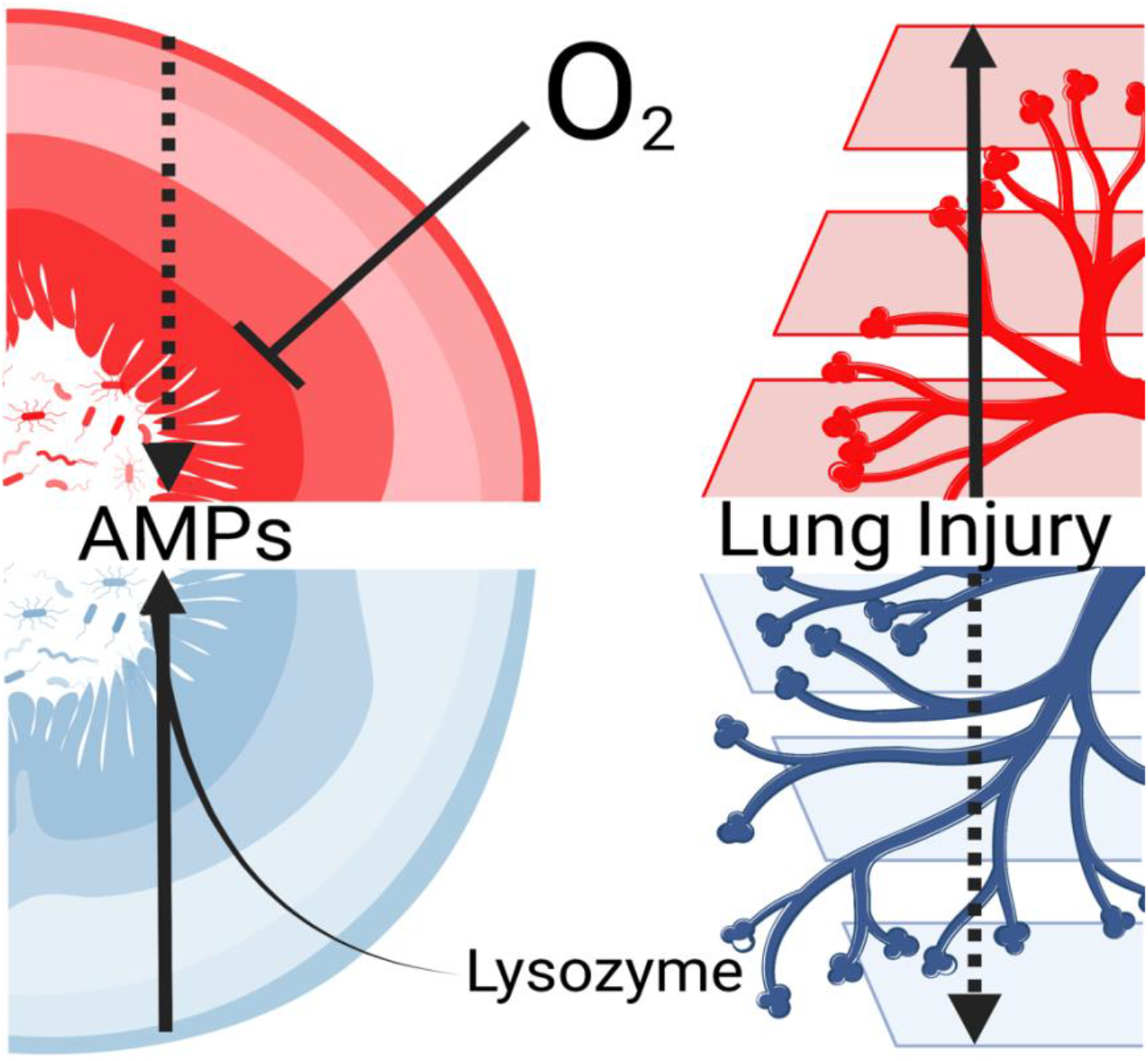

**In Brief:** Using a combination of murine models and organoids, Abdelgawad and Nicola et al. find that suppression of antimicrobial peptide release by the neonatal intestine in response to supra-physiological oxygen influences the progression of lung injury likely via modulation of the ileal microbiota.

**Highlights:**

- Supraphysiologic oxygen exposure alters intestinal antimicrobial peptides (AMPs).
- Intestinal AMP expression has an inverse relationship with the severity of lung injury.
- AMP-driven alterations in the intestinal microbiota form a gut-lung axis that modulates lung injury.
- AMPs may mediate a gut-lung axis that modulates lung injury.

## INTRODUCTION

Fetal lung development is a tightly regulated process normally occurring at low oxygen tension^1^. However, premature birth exposes the developing lung to far greater oxygen levels and this is usually compounded by the clinical use of supplemental oxygen and mechanical ventilation after birth^2^. The direct effects of supraphysiologic oxygen on the lungs are well described^3^, but there is also evidence for concurrent systemic, non-pulmonary oxygen toxicity exemplified by retinopathy of prematurity^3^. However, the effect of supraphysiologic oxygen on the host-microbiota interface of the neonatal intestine remains unclear.

Antimicrobial peptides (AMPs) are an extensive class of proteins secreted across mucosal surfaces that have central roles in the response to inflammation and the regulation of the commensal microbiota^4^. In addition to their antimicrobial properties, AMPs have anti-inflammatory, wound-healing, and tissue-protective effects^5^. In the newborn, intestinal and nasopharyngeal secretion of AMPs increases with age, potentially a necessary suppression to aid the establishment of a normal microbiota or to compensate for the intake of milk-derived AMPs^4^. Since AMP expression also increases in response to inflammation^5^, supraphysiologic oxygen might disrupt initial microbial colonization.

While exploring the effects of hyperoxia exposure on the transcriptional landscape of the developing gut and lungs, we were intrigued by alterations in AMP production in the neonatal intestine that were associated with alterations in the microbiota. Here, we asked if hyperoxia-induced alterations in intestinal AMP expression could alter the composition of the intestinal microbiota, creating a feedback loop that modulates inflammation in the developing lung.

## RESULTS

### Hyperoxia exposure alters lung morphology and function

To examine potential mechanistic connections between the effects of supraphysiological oxygen exposure on the intestine and on the lungs, we utilized a well-established model of bronchopulmonary dysplasia^6,7^ and exposed newborn mouse pups to hyperoxia (fraction of inspired oxygen, FiO_2_, 0.85), and their littermate controls to normoxia (FiO_2_ 0.21), from the third to the 14th day of life (P3-P14, Fig. 1A). We reproduced well-described hyperoxia-induced alterations in lung structure and function (Fig. 1B-C)^8^. We also observed changes in the pulmonary transcriptome and related regulatory pathways (Fig. S1) that are consistent with similar observations^9–12^ showing type 2 immune polarization and extensive alterations in alveolar epithelial, endothelial and fibroblast cell populations in response to hyperoxia-induced lung injury.

**Figure 1.**
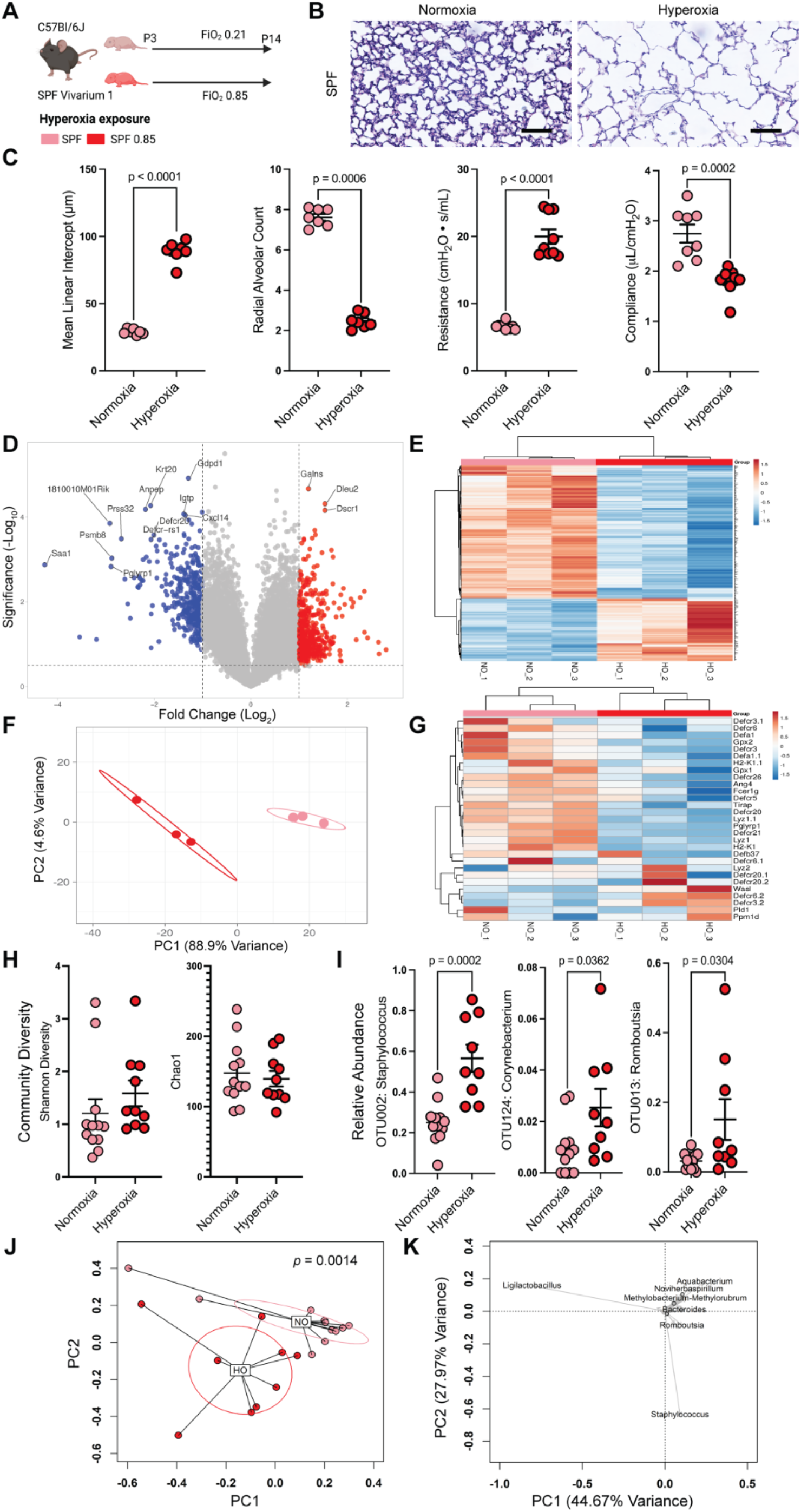
Oxygen exposure reduces intestinal antimicrobial peptide expression. A) Neonatal C57BL/6J mice were exposed to normoxia or hyperoxia from the 3^rd^-14^th^ day of life (n = 4 litters with 5-7 neonatal mice/litter per exposure group). FiO_2_, fraction of inspired oxygen. SPF, specific-pathogen-free. B) Representative photomicrographs of the distal lung sections of 14-day-old mice. C) Hyperoxia exposure is associated with alterations in lung morphology and function. Data are shown as mean ± SEM, with significance testing by two-tailed *t*-test. D) Volcano plot of ileal gene expression array showing gene expression altered by hyperoxia exposure. E) Heatmap showing genes regulated by hyperoxia exposure. F) Principal components analysis showing differential clustering of normoxia and hyperoxia exposed ileal genes. PC, principal component. G) Ileal antimicrobial peptide expression is decreased in hyperoxia-exposure mice. H) Community diversity of the adherent and luminal ileal bacterial microbiome is not significantly altered by hyperoxia exposure. I) The relative abundance of an operational taxonomic units (OTU 002) that aligns to the genus *Staphylococcus* increases after hyperoxia exposure, as do OTUs aligning to *Corynebacterium* (OTU 124) and *Romboutsia* (OTU 013). Data are shown as mean ± SEM, with significance testing by two-tailed *t*-test. J) Principal coordinates analysis of Bray-Curtis dissimilarity show global alterations in community composition in hyperoxia-exposed mice. Significance testing by permutational ANOVA (PERMANOVA), with equivocal dispersion confirmed by permutational multivariate analysis of dispersion (PERMDISP). PC, principal component. K) Loading plot of principal components analysis of Hellinger transformed Euclidian distances showing the contribution of specific genera to the global community composition. Schematic in (A) was generated using BioRender. See also Figures S1 and S2.

### Intestinal antimicrobial peptide expression is altered in hyperoxia-exposed neonatal mice

To interrogate potential hyperoxia-induced alterations in the neonatal gut, we then examined transcriptomic changes in the terminal ileum. Intriguingly, we found that hyperoxia-exposed mice had a significantly altered ileal transcriptome (Figure 1D-G). Global gene expression patterns were significantly altered (Figure 1D-F), including the prototypical AMP lysozyme (*Lyz1*, 4.2-fold decrease, FDR *p* = 0.012, Figure 1G, Table S1). In addition to other AMPs, additional markers of ileal Paneth cells were also decreased (*Pnliprp2*, 5.0-fold decrease; *Reg3g*, 2.7-fold decrease; and *Guca2b*, 2.4-fold decrease. Table S1). We verified these alterations in AMPs using qPCR (Figure S2A). Together, these results suggest that hyperoxia exposure reduces expression of intestinal AMPs.

### The intestinal microbiota responds to pulmonary hyperoxia exposure

We have reported that axenic (microbiota-free) mice devoid of a microbiome are protected^13^, while antibiotic-exposed mice with a disrupted microbiome are more susceptible, to hyperoxia-induced lung injury^7^. Together, these and other studies^14^, suggest that the intestinal microbiota may modulate hyperoxia-induced lung injury. Because of this evidence, and due to our observation of changes in the transcription of potentially microbiota-altering AMPs, we asked if hyperoxia exposure altered the intestinal microbiota.

To answer this question, we collected the intact terminal ileum from hyperoxia-exposed neonatal mice and littermate controls, which had been raised under specific pathogen-free (SPF) conditions and performed 16S rRNA MiSeq to define both the adherent and luminal ileal microbiome. We found the global community composition but not diversity of the ileal microbiota was significantly altered by exposure to hyperoxia (*p* = 0.0014, pseudo-*F* = 6.3401, permutational ANOVA (PERMANOVA) of Bray-Curtis dissimilarity, *p* = 1, permutational multivariate analysis of dispersion (PERMDISP), Figure 1H-K). The first and second principal components are driven primarily by the abundance of the genus *Staphylococcus* and several uncultured *Ligilactobacillus* species (Figure 1J). In unadjusted univariate analysis, we identified increases in taxa aligning to the genera *Staphylococcus, Corynebacterium* and *Romboutsia* (Figure 1I). We verified these alterations using negative binomial regression (Figure S2), which also detected significant alterations in *Proteus* and Lachnospiraceae NK4A136 group. Correlation analysis between these differentially abundant genera and our validation qPCR for key AMPs supports a direct link between the expression of *Lyz1* and the relative abundance of *Staphylococcus* (Figure S2F). Together, these results suggest hyperoxia exposure alters the ileal microbiome by increasing the expression of lysozyme which is associated with an increase in the relative abundance of *Staphylococcus*.

### Hyperoxia directly suppresses intestinal epithelial AMP expression

We turned to small intestinal organoids to directly test the capacity for hyperoxia exposure to depress intestinal AMP expression. We exposed organoids to either hyperoxia (FiO_2_ 0.95) or normoxia (FiO_2_ 0.21) for 24 hours (Figure 2A). In general, live imaging with visual light microscopy and immunohistochemistry demonstrated organoids maintained similar shape and cellular composition in both normoxia and hyperoxia (Figure 2B-C, Figure S3), but with a decrease in epithelial integrity (*p* < 0.0001, two-tailed *t*-test, Figure 2B). RNAseq revealed transcriptomic changes that generally recapitulated the changes observed in hyperoxia-exposed mice (Figure 2D-G). Principal components analysis shows a clear separation in gene expression between hyperoxia and normoxia-exposed organoids (Figure 2D), which is also reflected in a heatmap of regulated genes (Figure 2E). Pathway analysis showed extensive upregulation of genes associated with cellular inflammatory response and response to injury (Figure 2G). Like *in vivo*, we also noted a decrease in several Paneth cell markers (Table S2). We verified the relative expression of several genes by qPCR. As indicated in our transcriptomic data, *Lyz1* was decreased in mice exposed to hyperoxia, as was the defensin *Defa2* and the Paneth cell marker *Pnliprp2* (Figure S2A). Together these data suggest that the downregulation of AMP expression is linked to cellular damage response of the ileal epithelium from exposure to supraphysiologic oxygen.

**Figure 2.**
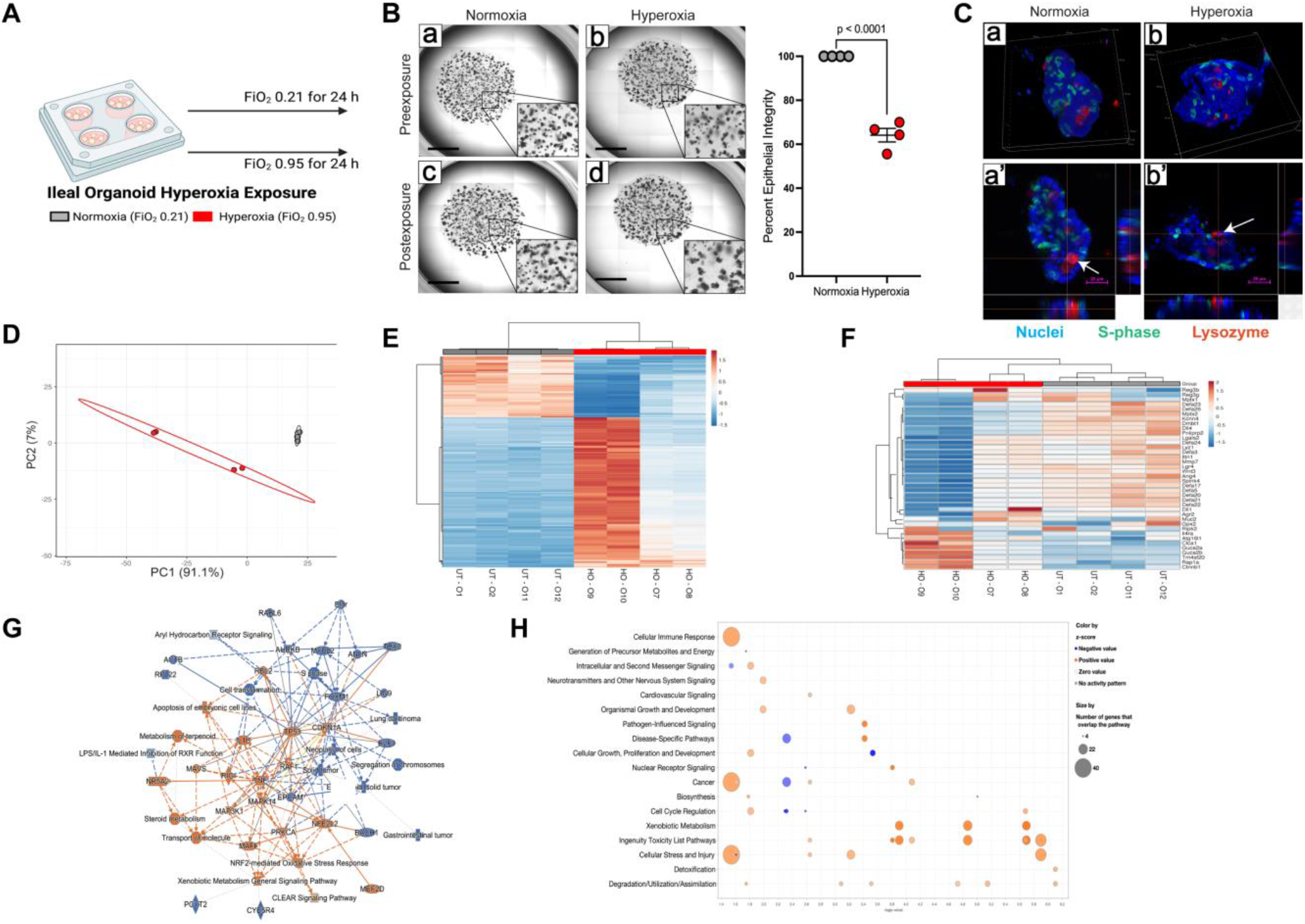
Antimicrobial peptide expression is reduced in hyperoxia-exposed intestinal organoids. A) Small intestinal spheroid organoids derived from neonatal C57BL/6J mice were exposed to either hyperoxia or normoxia for 24 hours. FiO_2_, fraction of inspired oxygen. B) Representative images of organoids before and after exposure, with insets at 40x magnification. The percentage of organoids wi healthy appearing epithelium declined in hyperoxia-exposed organoids. Data are shown as mean ± SEM, with significance testing by a two-tailed *t*-test. Scale bar represents 1000 µm. C) Representative immunohistochemistry after exposure to normoxia or hyperoxia. Nuclei in blue, actively proliferating cells in gree and lysozyme-positive cells in red. Arrows identify lysozyme-positive Paneth cells. Scale bar represents 25 µm. D) Principal components analysis showing differential clustering of normoxia and hyperoxia exposed ileal genes. PC, principal component. E) Heatmap showing genes regulated by hyperoxia exposure. F) Heatmap of antimicrobial peptide expression is decreased in hyperoxia-exposure organoids. G) Ingenuity pathway analysis showing regulated pathways in hyperoxia or normoxia. H) Bubble plot showing up and down-regulated pathways from hyperoxia exposure. The schematic in (A) was generated using BioRender.

### Augmentation of lysozyme improves lung function in hyperoxia-exposed mice

Having shown that exposure to supraphysiologic oxygen resulted in depressed expression of intestinal AMPs and an altered microbiome, we next wanted to test if AMP expression drove this effect. Lysozyme is the principal AMP, and expression is reduced in early life on most mucosal surfaces^15^. To test if the suppression of AMP expression in hyperoxia-exposed mice might have functional consequences, we repeated the hyperoxia-exposure experiment above using nearly genetically identical mice (C57BL/6NCrl, Figure 3A). However, to naturally vary their microbiota as much as possible, we purchased these animals from a different vendor and raised them in a different SPF vivarium. We exposed a randomly selected subset of their neonates to lysozyme by oral gavage and compared them to unexposed littermates. Surprisingly, augmenting intestinal lysozyme improved lung structure and function after hyperoxia exposure (MLI *p* < 0.0001, RAC *p* < 0.0001, and resistance *p =* 0.0212, two-way ANOVA, Figure 3B-C).

**Figure 3.**
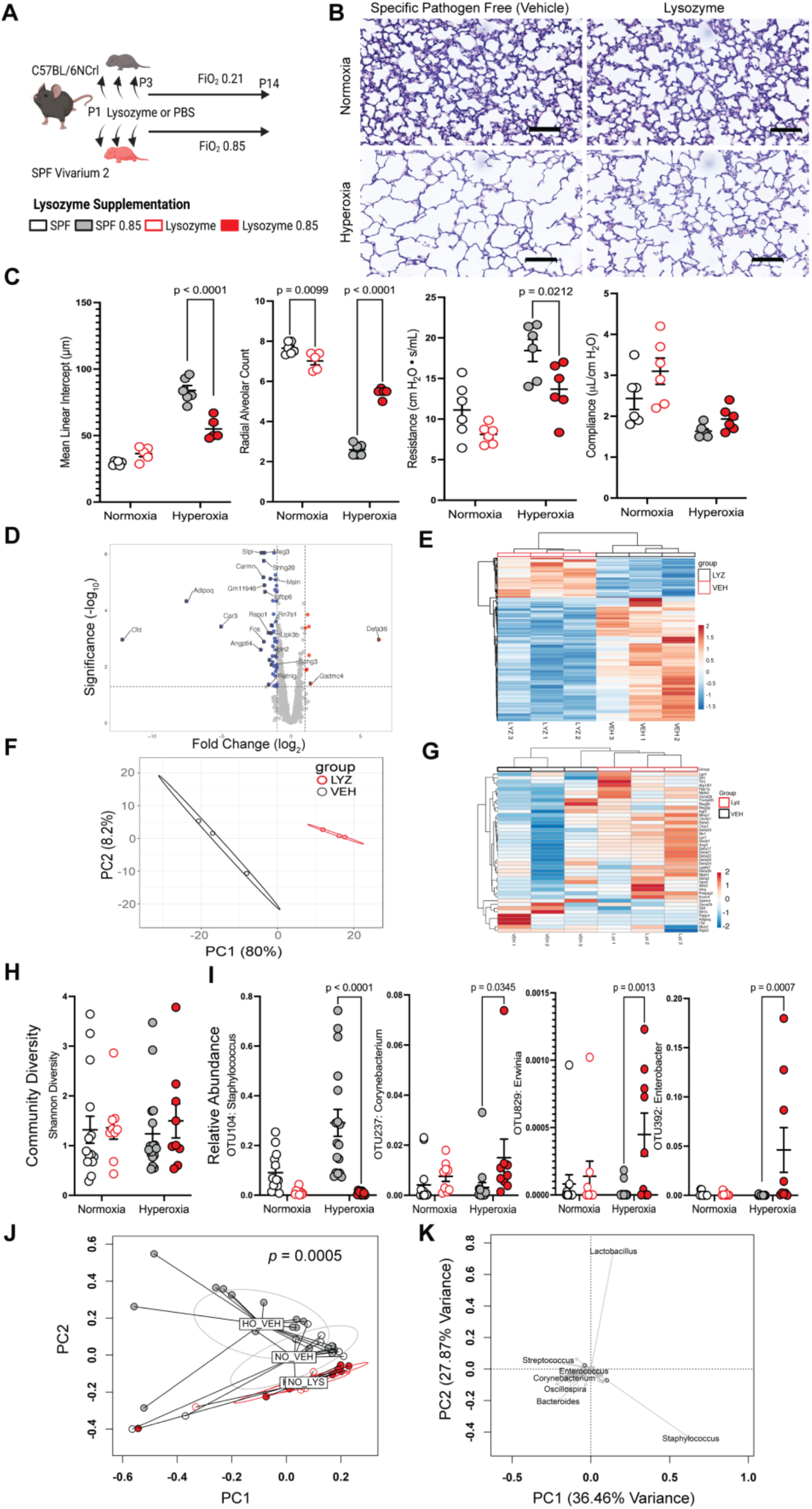
Intestinal lysozyme supplementation reduces hyperoxia-induced lung injury. A) Neonatal C57BL/6NCrl mice randomized to either every other day exposure to lysozyme by gastric gavage or their littermate controls were then exposed to normoxia or hyperoxia from the 3^rd^-14^th^ day of life (n = 4 litters with 5-7 neonatal mice/litter per exposure group). FiO_2_, fraction of inspired oxygen. PBS, phosphate-buffered saline (vehicle). SPF, specific-pathogen-free. B) Representative photomicrographs of the distal lung sections of 14-day-old mice. C) Lysozyme exposure ameliorates hyperoxia-induced disruptions in lung morphology and function. Data are shown as mean ± SEM, with significance testing by two-way ANOVA. D) Volcano plot of ileal RNAseq showing gene expression altered by lysozyme exposure. E) Heatmap showing genes regulated by lysozyme exposure. F) Principal components analysis showing differential clustering of ileal genes in lysozyme exposed mice. PC, principal component. G) Ileal antimicrobial peptide expression is altered in lysozyme-exposed mice. H) The community diversity of the adherent and luminal ileal bacterial microbiome is not significantly altered by lysozyme exposure. I) The hyperoxia-induced increase in the relative abundance of operational taxonomic unit 014 (*Staphylococcus*) is ameliorated by lysozyme exposure. Multiple other genera are increased in lysozyme and hyperoxia-exposed mice. Data are shown as mean ± SEM, with significance testing by two-way ANOVA. J) Principal coordinates analysis of Bray-Curtis dissimilarity show global alterations in community composition in lysozyme exposed mice. Significance testing by permutational ANOVA (PERMANOVA), with equivocal dispersion confirmed by permutational multivariate analysis of dispersion (PERMDISP). PC, principal component. K) Loading plot of a principal components analysis of a Hellinger transformed Euclidian distance showing global community composition significantly altered in lysozyme-exposed mice. The schematic in (A) was generated using BioRender. See also Figure S3.

### Lysozyme augmentation increases intestinal AMP expression

To identify alterations in intestinal AMP expression resulting from lysozyme exposure, we performed RNAseq using the terminal ileum of normoxia exposed mice. Intriguingly, we found that lysozyme-exposed mice had a significantly altered ileal transcriptome (Figure 3D-G). Global gene expression patterns were significantly altered (Figure 3D-F), including altered expression of multiple AMPs (Figure 3G). Of note, *Lyz1* expression increased 2.433-fold (FDR *p* = 0.0004).

Together, these results suggest that dietary lysozyme exposure alters expression of the ileal transcriptome and results in additional ileal lysozyme production in an apparent feed-forward mechanism.

### Lysozyme alters the intestinal microbiota

We next asked if supplementary lysozyme altered the intestinal microbiota. Oral lysozyme supplementation produced a marked change in the overall composition of the ileal microbiota as compared to littermate controls (*p* < 0.0001, pseudo*-F* = 0.2544, PERMANOVA; *p =* 0.7824, PERMDISP, Figures 3J and S4), without significantly altering the community diversity (Figure 3H). On univariate analysis, multiple species were differentially abundant (Figure 3I). Most intriguing, the robust increase of *Staphylococcus* in hyperoxia-exposed Jackson Laboratory mice, was replicated in the vehicle and hyperoxia-exposed SPF Charles River mice but suppressed in lysozyme-exposed animals (*p* < 0.0001, two-way ANOVA, Figure 3I). Robust increases were also observed in mice exposed to both lysozyme and hyperoxia. *Corynebacterium, Erwinia*, and *Enterobacter* also increased in these doubly exposed mice (*p =* 0.0345, *p =* 0.013, *p =* 0.0007, two-way ANOVA, Figure 3H). MaAsLin 2 and binomial regression identified a shared 48 OTUs altered by lysozyme exposure (Figure S4) in addition to these four genera. Together, these analyses suggest that augmenting dietary lysozyme results in a robust alteration of the ileal microbiome and ameliorates the increase in *Staphylococcus* that otherwise occurs with hyperoxia exposure.

### Lysozyme augmentation alters the pulmonary transcriptome

We next sought to examine if the functional improvements we identified in the lungs of mice exposed to lysozyme were associated with differences in the lung transcriptome. Indeed, RNAseq revealed extensive alterations in gene expression in the lung (Figure 4). While the most increased inflammatory related genes remained elevated in lysozyme and hyperoxia-exposed mice they were reduced as compared to controls exposed only to hyperoxia (Table S3). Similar patterns exist for genes related to endothelial regulation (Table S4). However, multiple genes exhibited opposite regulation in lysozyme-exposed mice (Table S5). Overall, the most reduced genes in hyperoxia exposure, were reduced regardless of lysozyme exposure (Figure 4A-B). The genes with the most increased expression were similar (Figure 4A-B). However, notable differences in *Hpgd, Akr1b8, Timp1* and *Zmat3* were appreciable. Altogether, 432 genes were differentially regulated in vehicle-exposed controls (Figure 4C) of which 271 were also differentially regulated the lysozyme exposed mice (Figure 4D), leaving 149 unique differentially regulated genes in lysozyme exposed mice (Figure 4E). The most significantly upregulated pathways in vehicle-exposed controls, included fibrosis, wound healing and IL-6-type cytokine signaling-related pathways (Figure 4F).

**Figure 4.**
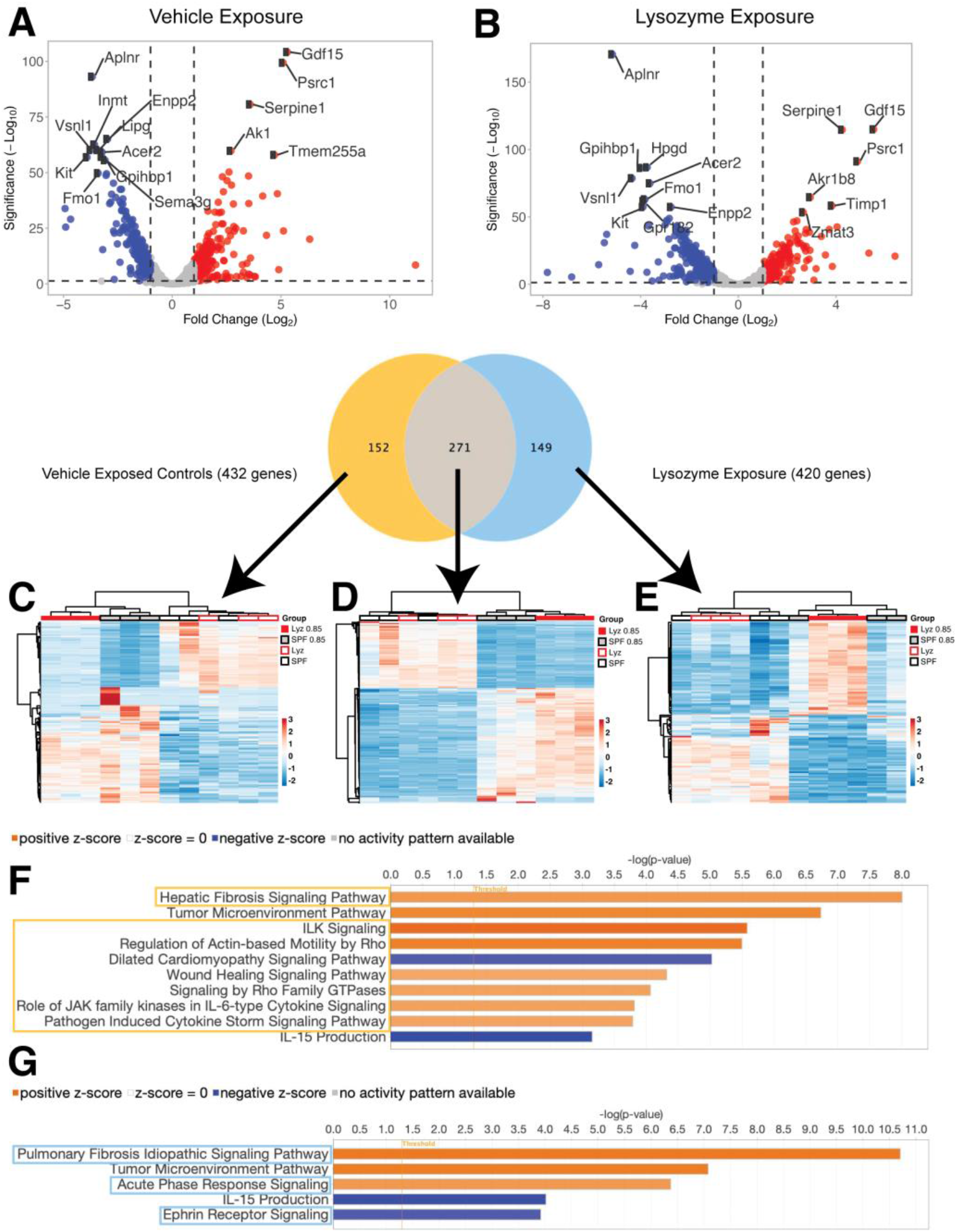
Lysozyme exposure alters the lung transcriptome. A) Volcano plot showing hyperoxia alters gene expression in vehicle-exposed controls. B) Volcano plot showing lysozyme exposure alters gene expression in the lung. C) Heatmap showing differentially expressed genes in vehicle-exposed controls. D) Heatmap showing similarly expressed genes between all groups. E) Heatmap showing differentially expressed genes in lysozyme-exposed mice. F) Major pathways altered in mice only exposed to normoxia or hyperoxia. G) Major pathways altered in lysozyme-exposed mice.

However, different fibrosis-related and acute phase response pathways were upregulated in lysozyme-exposed mice (Figure 4F). Together, these transcriptional differences characterize a unique response to hyperoxia in the lungs of mice exposed to lysozyme, that is associated with ameliorated injury and an improvement in lung structure and function.

## DISCUSSION

Here, we have shown that exposure to supraphysiologic oxygen alters intestinal AMP expression, creating a microbiota-mediated response that modulates lung injury. Hyperoxia inhibited AMP transcription, both *in vivo* in neonatal mice and *in vitro* in intestinal organoids. *In vivo*, this altered the intestinal microbiota. Conversely, augmenting AMP expression altered the intestinal microbiota and improved lung injury. This reciprocal relationship between intestinal AMPs and lung injury demonstrates a gut-lung axis mechanism that can influence hyperoxia-exposure-induced lung injury. Together, these findings suggest intestinal AMPs are potential modulators of lung injury and repair.

AMPs are small cationic peptides that play a key role in host defense against microbes^16^. AMPs have broad antimicrobial activity and can kill or inhibit the growth of bacteria, fungi, and viruses^17^. AMPs, therefore, also play a key role in the regulation of the intestinal microbiota^16^. However, the potential bioactive effects of AMPs are not exclusively related to manipulating the microbiota. Lysozyme, the major AMP, can directly reduce the propagation of reactive oxygen species in colitis by releasing bacterial superoxide dismutase from *Lactobacillus lactis*^*18*^. Numerous species of bacteria within the microbiota contain superoxide dismutase genes^18^. This suggests potential mechanisms by which the redox balance of the intestinal epithelium might be influenced by alterations in AMP expression. Multiple potential links between the lung and the gut have been proposed to explain the gut-lung axis^21^.

Samuelson and colleagues^22^ have proposed that microbes or microbial components may be trafficked to the lung via the lymph. Similarly, metabolites produced or modified by the intestinal microbiota, such as small-chain fatty acids, may also influence lung physiology^23^. Lysozyme can also liberate ligands for nucleotide binding oligomerization domain containing 1 (Nod1) from bacterial cell wall peptidoglycans. Circulating Nod1 ligands may reach the lung directly or may participate in immune cell maturation in the gut to determine the set points of the innate immune system in early life^24,25^, and to educate the adaptive immune system^26^. The AMP-mediated gut-lung axis we describe here could impact some or all these potential pathways.

Exposure to supplemental oxygen is one of the most ubiquitous clinical interventions. Preterm infants, particularly the most immature, often require exposure for prolonged periods^2^. The disastrous effects of such exposure on the structure and function of neonatal lungs are well described^3^. In the adult gut, the gradient between hypoxia to anoxia across the radial axis of the gut has a profound influence on the composition of the intestinal microbiota and contributes to the maintenance of distinct communities of adherent and luminal microbes^27,28^. Hyperbaric oxygen exposure can shift the composition of these mature communities^27,28^, as can inflammation, which alters the oxygen gradient in enterocytes^*29*^. However, the maturation of intestinal microbial communities is a developmental process, and initially, during the newborn period, the gut lumen is less hypoxic^30^.

Recent evidence suggests that this process is driven by the accumulation of microbial biomass in the more distal intestine^30^. This is supported by the developmental progression from aerobic to facultative anaerobic microbes in the neonatal gut, and by the observation that while the intestinal lumen of axenic mice is anaerobic this appears to be driven by a slower chemical process^30^. This suggests our findings may be influenced by a unique developmental window created by the immature gut being more vulnerable to oxidative stress and the immature microbiota being less capable of functioning as an anoxic sink.

In adult mice, hyperoxia exposure has been shown to alter the lung and intestinal microbiota. Ashley and colleagues observed an increase in the oxygen-tolerant genera *Staphylococcus* after 72 hours of exposure to supraphysiological oxygen^14^. We observed a similar increase in *Staphylococcus* in the ileal microbiota of neonatal mice exposed to hyperoxia that replicated robustly in mice with different starting microbiomes. While Ashley and colleagues confirmed a previously described^29^ decrease in the Ruminococcaceae after hyperoxia exposure in the cecal microbiota of these adult mice^14^, several differences likely explain our divergent findings. First, we sampled the ileal microbiota, which is more translationally relevant to neonatal intestinal pathology. Second, our neonatal mice were considerably smaller and exposed to hyperoxia for a longer duration, which would be intolerable for adult mice. Finally, their intestinal microbiota were in an earlier stage of community formation, which places their microbial communities under considerably different ecological pressures.

Intriguingly, this increase in *Staphylococcus* was ameliorated in our lysozyme-exposed and associated with less severe lung injury. In subsequent experiments, Ashley and colleagues further found that adult axenic mice were protected from lung injury, and antibiotic-exposed mice experienced worse lung injury^14^ - findings we have previously demonstrated in neonatal mice^7,13^.

Under SPF conditions, the intestinal microbiota of co-housed mice converges over time^31,32^, and the primary source of the intestinal microbiota is almost universally the mother but may show some littermate effects^33,34^. We took multiple steps to address these factors in these experiments. First, we intentionally varied the initial microbiota of the maternal mice as much as possible by purchasing them from two different animal vendors, cage-randomizing them for cohousing on arrival, and acclimatizing them in two different SPF vivaria before breeding. Second, to limit a littermate effect, we took the newborn mice from two simultaneously born litters and randomly redistributed them between their two mothers on the day of birth. We then assigned one pooled litter to hyperoxia and the other to normoxia and rotated the mothers daily to minimize their unique contributions. For exposure to lysozyme, we randomly selected newborn mice within these homogenized litters and used their unexposed littermates as controls. Because this approach likely underestimates the potential microbiota effects of lysozyme exposure due to littermate cross-contamination, it underscores that increase in *Staphylococcus* abundance with hyperoxia exposure is robust, since we replicated it across a wide range of initial microbiomes.

In summary, we have described a gut-lung axis driven by intestinal AMP expression and mediated by the intestinal microbiota that influences hyperoxia-induced lung injury. These murine and organoid experiments suggest that AMP expression represents a potential therapeutic target to modulate the intestinal microbiota and the response to lung injury. These results have implications for the clinical management of premature infants in neonatal care at high risk of developing bronchopulmonary dysplasia.

## METHODS SUMMARY

Detailed methods are provided below and include the following:

- KEY RESOURCES TABLE
- RESOURCE AVAILABILITY
  - Lead contact
  - Materials availability
  - Data and code availability
- EXPERIMENTAL MODEL AND SUBJECT DETAILS
  - Animals
  - Mouse intestinal spheroid organoids
- METHOD DETAILS
  - FlexiVent pulmonary function
  - Histology and morphometry
  - Immunohistochemistry
  - Generation of intestinal organoids
  - Organoid hyperoxia experiment
  - Transcriptomics
  - Microbiome analysis
  - Bioinformatics
  - Statistical analysis

## Acknowledgments

Research reported in this article was supported by the NIH: K08 HL151907 (KW), K08 HL141652 (CL), K08 DK120871 (AO), 1U01ES027697 (TJ) and R21ES031559 (TJ); the Kaul Pediatric Research Institute at Children’s of Alabama (KW); the Microbiome Center at UAB (KW); and the Harvard Digestive Disease Center (AO). The authors thank Joseph Pierre for his advice.

## Author contributions

Conceptualization: NA, AO, TJ, KW. Experimentation: AA, TN, BH, KT, CA, PJ, CR, AO, TJ, KW. Analysis: IM, NA, AO, TJ, KW. Supervision: AO, TJ, KW. Manuscript: KW. All authors approved of the final submitted version of the manuscript.

## Inclusion and Diversity

We support the inclusive, diverse, and equitable conduct of research. We worked to ensure sex balance in the selection of non-human subjects. Multiple authors of this paper self-identify as an underrepresented ethnic minority in their field of research, as a gender minority, or as a member of the LGBTQA+ community. While citing references scientifically relevant to this work, we also actively worked to promote gender balance in our reference list.

## METHOD DETAILS

### Key Resource Table

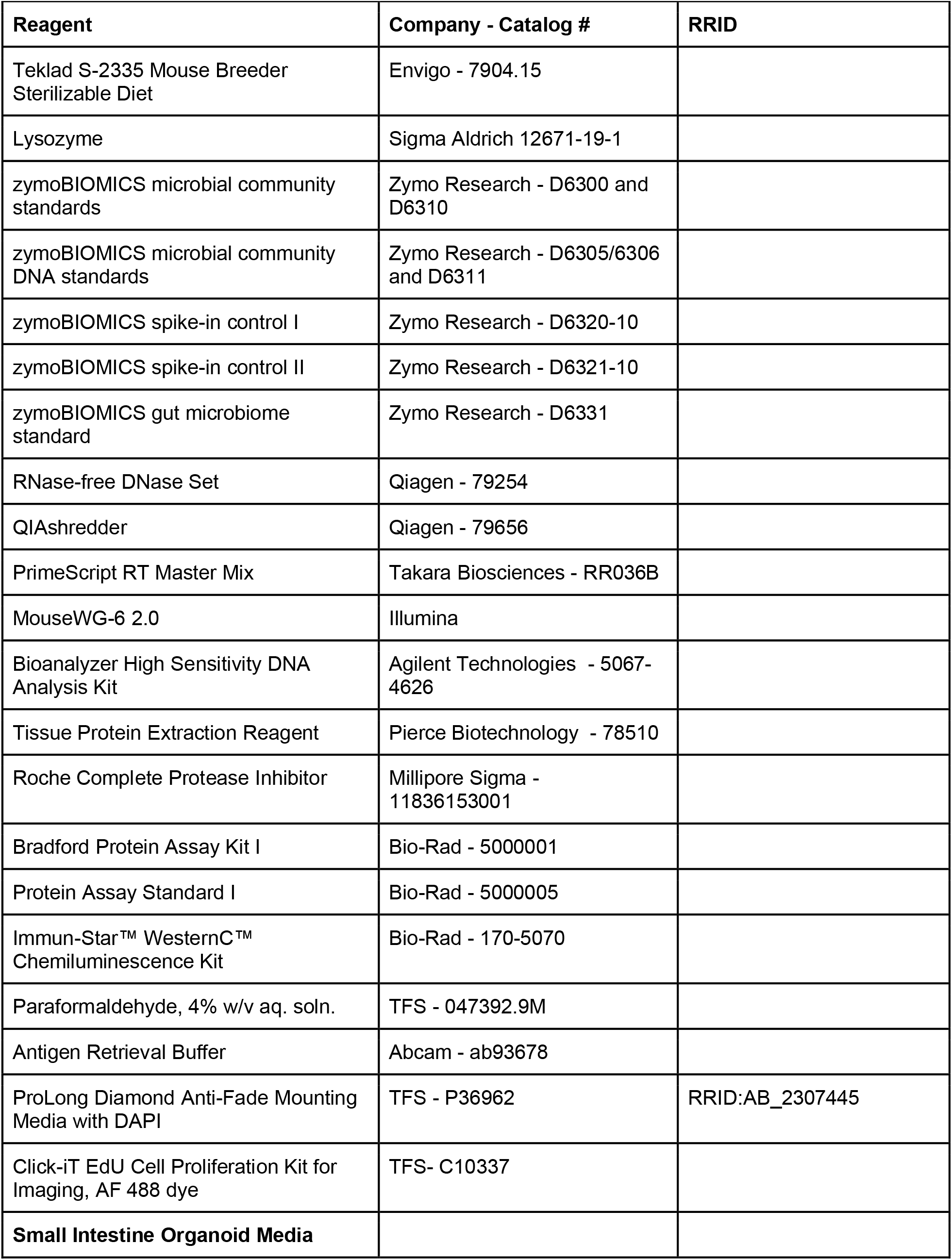
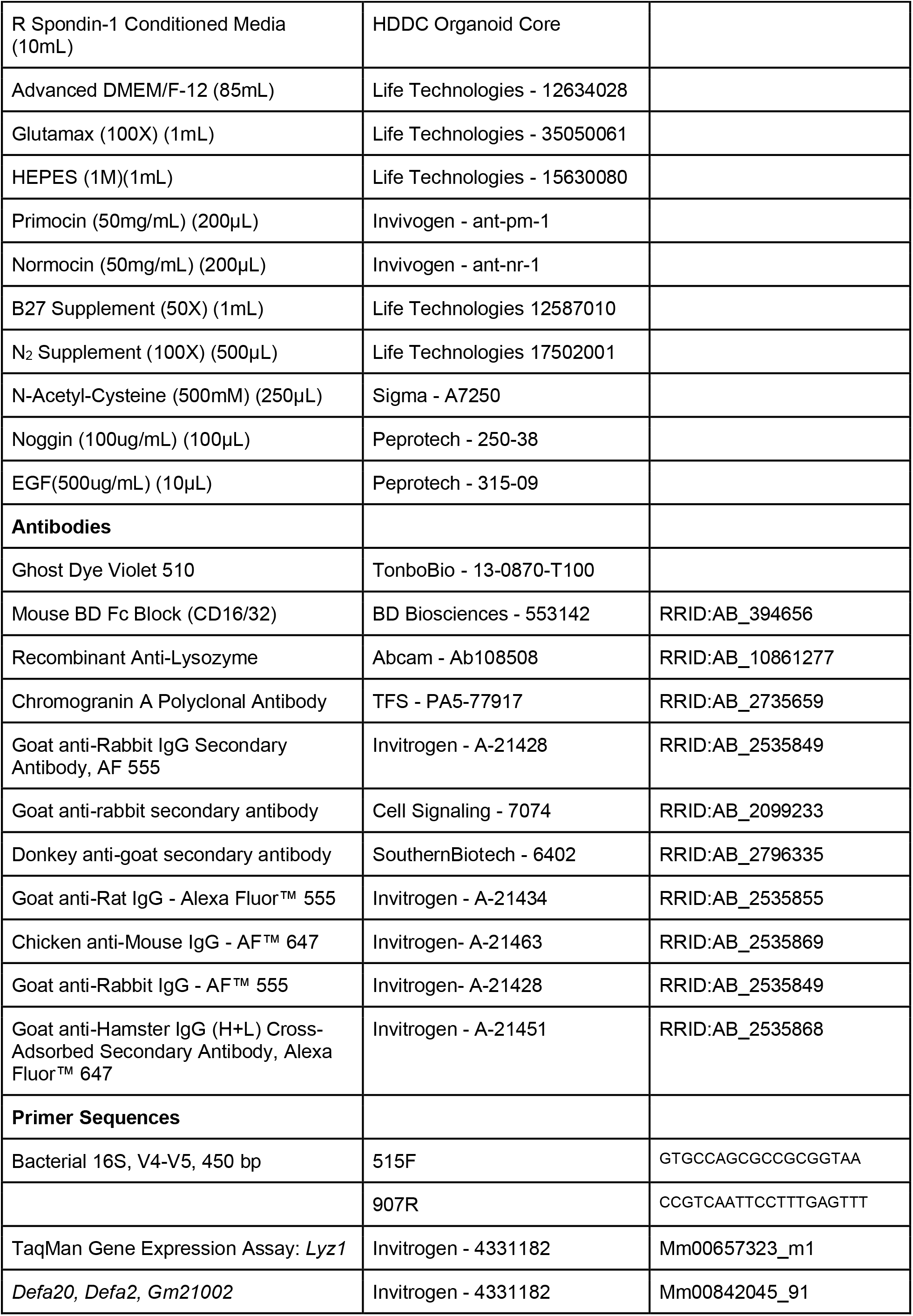
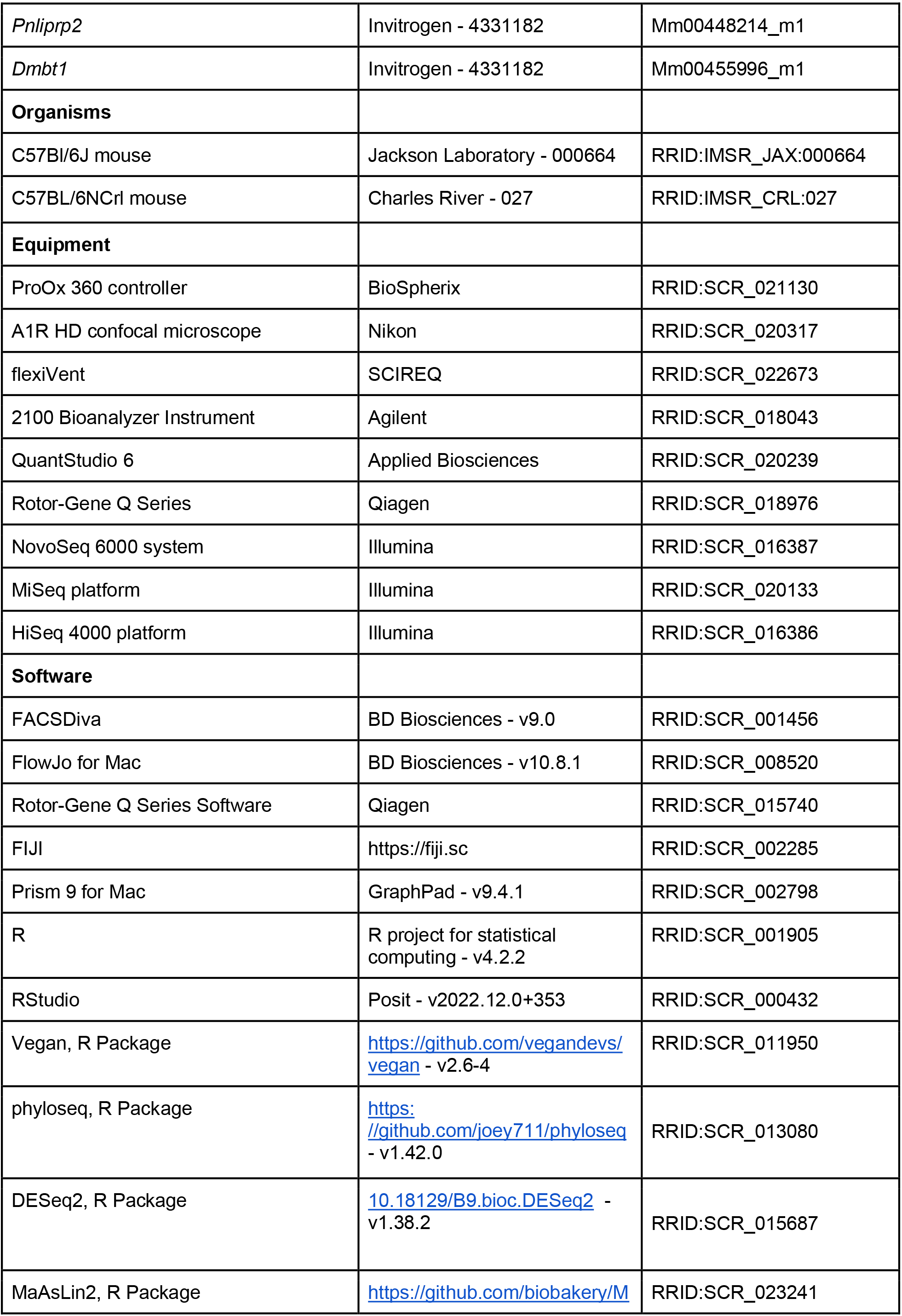
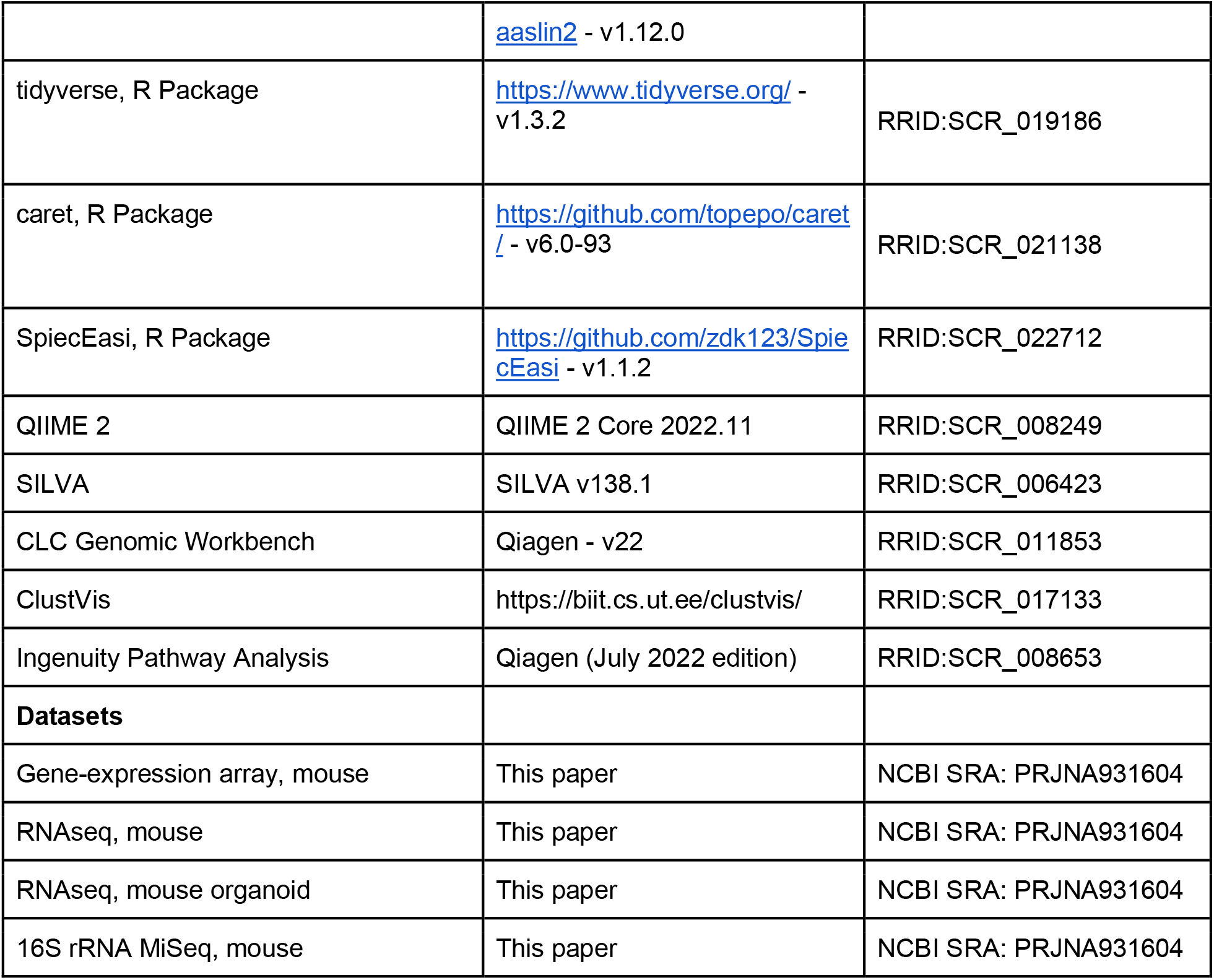

### Resource Availability

#### Lead Contact

Further information and requests for resources and reagents should be directed to and will be fulfilled by the lead contact, Dr. Kent Willis.

#### Materials Availability

This study did not generate any unique reagents or transgenic animals.

#### Data Availability

- 16S and RNA sequencing data were deposited at the NCBI Sequence Read Archive (BioProject ID: PRJNA931604).
- Gene expression array data were deposited at the NCBI Gene Expression Omnibus (Accession number: GSE125489).
- Processed data files are available at github.com/WillisLungLab/AMP.
- This manuscript does not report original code.
- All other data needed to evaluate the conclusions in the manuscript are available within the main text or supplementary materials.

### Experimental Model Details

#### Animals

Animal experimentation was performed at the University of Alabama at Birmingham. To broaden the translatability and reproducibility of the resulting experiments, we intentionally selected two similar, but not identical, mouse strains that we purchased from different animal vendors and raised in different vivaria to vary their initial microbiomes. In the first experiment, we used 8-week-old female C57Bl/6J mice (Stock Number 000664) purchased from the Jackson Laboratory (hence, JAX) and allowed to acclimate in specific-pathogen free (SPF) housing with 4 animals per cage, on a 12-hour light/dark cycle, at 20-24C, with *ad libitum* access to irradiated rodent chow (Teklad 7904, Envigo) in SPF Vivarium 1 (Pittman Biomedical Research Building II) for two weeks. Similarly, for the second experiment, we purchased C57Bl/6NCrl mice (Stock Number 027) from the Charles River Laboratory and acclimated them in SPF Vivarium 2 (Volker Hall) with the same diet for two weeks. Animal studies were performed in accordance with the recommendations in the Guide for the Care and Use of Laboratory Animals of the National Institutes of Health, under protocol 22042 approved by the Institutional Animal Care and Use Committee at UAB. Animal experimentation is reported in accordance with the ARRIVE guidelines^1^. We utilized our established, standardized hyperoxia-exposure model of BPD^2,3^. We time-mated the mice to produce multiple simultaneous birth cohorts with the same perinatal exposure. On the day of birth, pups from two consecutively born litters with the same perinatal exposure were pooled and the pups were re-distributed evenly between the two dams. Litter sizes for all experiments were adjusted to 5–8 pups per treatment group to minimize nutritional effects on lung development. One pooled litter was randomly assigned to hyperoxia [fraction of inspired oxygen (FiO_2_) 0.85]) and the other to air (FiO_2_ 0.21). Pups were exposed continuously for 11 days, starting on the third day after birth until the 14th day (P3-14).

Oxygen concentrations were maintained using a ProOx monitor (Biospherix). Dams were rotated daily between the two pooled litters to limit maternal complications of hyperoxia. Growth was monitored daily for all pups throughout the experiment.

For the lysozyme exposure experiment, pups from two pooled litters at time were randomized on P2 to either lysozyme exposure or vehicle exposure. Lysozyme exposure was performed via every other day oral gavage of 10,000 units/gram in PBS to pups marked by tail clipping. Unmarked littermates were used as vehicle (PBS)-exposed controls. Experiments were repeated at least three times.

#### Mouse intestinal epithelial spheroid organoids

Organoid modeling was performed at the Harvard Digestive Disease Center (HDDC) Organoid Core at Boston Children’s Hospital. Crypts were collected from C57Bl6J mice and cultured in 50mL Matrigel and fed with 500mL murine enteroid media in a 24 well plate in a 37°C humidified incubator with 5% CO_2_.

Media was changed every 2-3 days for the duration of the cultures. Organoids were passaged every 7-10 days by dissolving the Matrigel in Cell Recovery Media with mechanical disruption.

### Further Method Details

#### Forced oscillometry

A subset of P14 mice not used for morphometry or transcriptomics were sedated with ketamine and xylazine via I.P. injection, and the flexiVent system (SCIREQ) was used to assess respiratory resistance and compliance as described(15, 17). Briefly, the mice were tracheotomized with the appropriately sized cannula secured with a silk ligature. The flexiVent then executed the neonatal mouse pulmonary function program using room air in the sedated, closed-chest animal. Calibration of the flexiVent was done using the tracheal cannula before each experiment. Lung volumes were measured by volume displacement after completion of the flexiVent measurements.

#### Histology and morphometry

The right lung was gravity inflated to 20 cmH_2_O via tracheal insertion of an angiocath with 4% formalin as described(7). Three random 4-5 µm sections from each lung were stained with hematoxylin and eosin.

To perform morphometry, lung sections were digitized under x20 magnification and analyzed independently by two researchers blinded to the group assignments using established methods (7, 18– 20). Alveolarization was quantified using the mean linear intercept (*L*_*m*_) and radial alveolar counts (RAC).

#### Immunohistochemistry

Organoids were kept in 1.5 ml Eppendorf tubes. Three tubes per group were fixed by 0.4% PFA. Samples were resuspended using a Wide Bore Pipet Tips in a blocking buffer (1% BSA, 1% normal donkey serum, 1% CD16/32 in PBS) for 30 min in RT, and then incubated for one hour with primary antibodies at RT in incubation buffer (1% BSA, 1% normal donkey serum, 0.3% Triton X-100, and 0.01% sodium azide in PBS). The samples then were washed in PBS, and then incubated for one hour with the corresponding secondary antibodies at RT in the incubation buffer. Samples were centrifuged and resuspended in the proper buffer for each step. Finally, samples were counterstained with Click-iT™ EdU Cell Proliferation Kit for Imaging, Alexa Fluor™ 488. Images were then captured using an A1R HD inverted confocal microscope.

#### Generation of intestinal organoids

Organoids were generated from C57Bl/6 mice after euthanasia using CO_2_ followed by cervical dislocation. Proximal duodenal tissue was dissected, and feculent matter was cleared using cold intraluminal PBS. The tissue was cut longitudinally and then cut into 0.5 cm pieces and cleaned repeatedly in PBS until the supernatant was cleared. The tissue was then placed in 2mM EDTA and incubated on ice with rocking for 30 minutes. The tissue was then shaken vigorously for 2 minutes to release crypts and pipetted up and down 25 times with a 10mL serological pipet while mixing simultaneously. The solution was passed through a 70uM filter, and the filtrate was collected into a new 50mL conical tube. The filtrate was centrifuged (5min at 300xg) and supernatant discarded. Dissociated cells were then washed three times in Advanced DMEM/F12, centrifuged (5min at 300xg) after each wash, and reconstituted in Matrigel (50uL/well) and plated in 24-well plates and incubated for 10 minutes at 37 C. Next, 0.5mL growth media was added to the wells. Enteroids were fed with growth media every 2-3 days and passaged approximately every 7 days.

### Murine Small Intestine (ENR) Media (100mL)

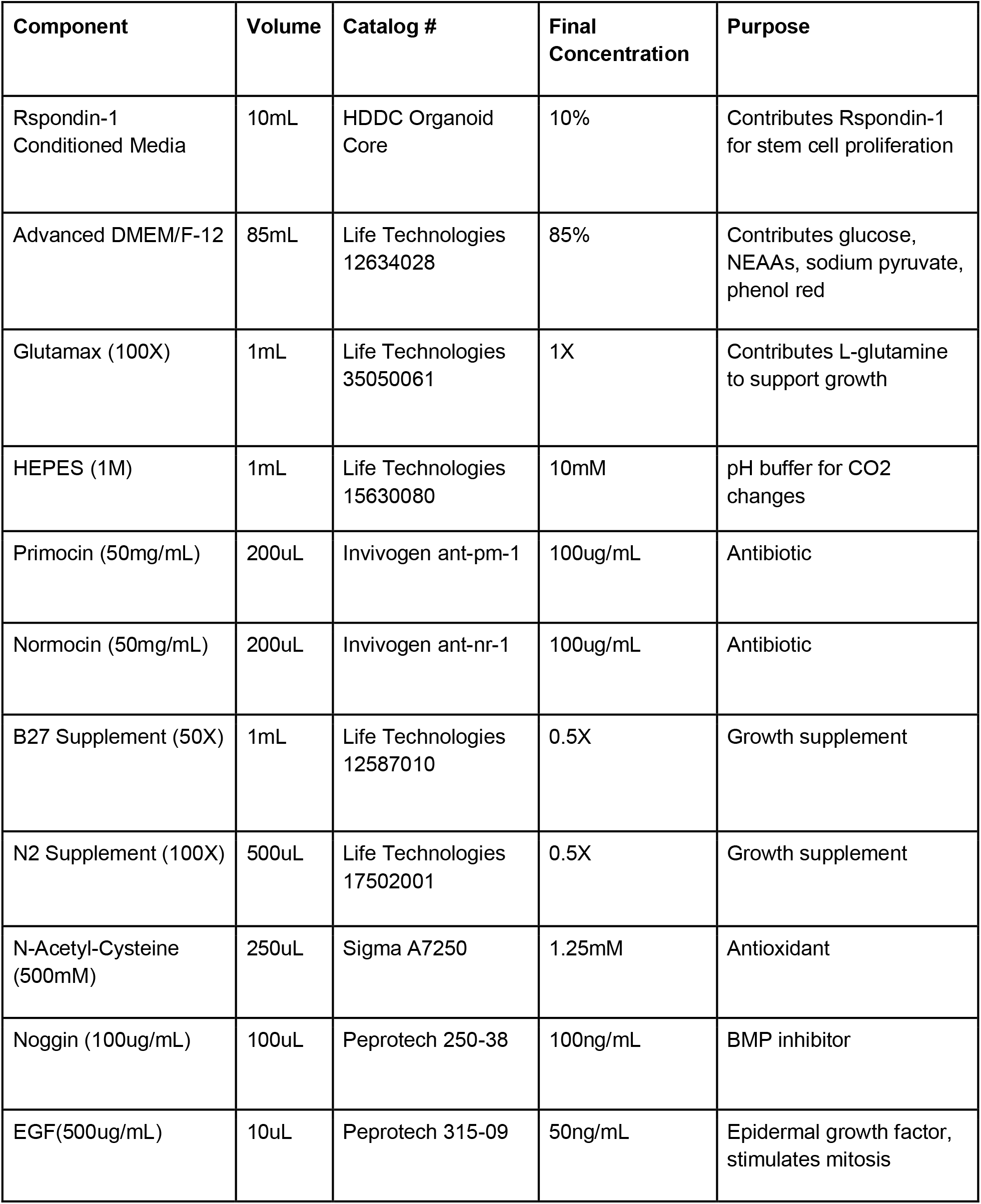

#### Organoid hyperoxia experiment

Organoid hyperoxia experiment. Intestinal organoids were generated as above and incubated in a hyperoxia chamber (StemCell Technologies) at 95% oxygen for 24 hours. After the hyperoxia was completed, the supernatant was removed and the organoids and Matrigel were collected in 0.5mL cell recovery media (Corning) per well and placed on ice for 40 minutes. The solution was then centrifuged (5 minutes at 300 g at 4C) and the supernatant was removed. The organoids were then either placed in 2% β-mercaptoethanol in RLT buffer or 4% paraformaldehyde for RNA isolation or fixation, respectively.

#### Transcriptomics

25 mg tissue sections from either the left lower lobe of the lung or from the terminal ileum were bead-lysed in RLT buffer (Qiagen) supplemented with 1% (w/w) 2-mercaptoethanol and frozen. RNA was isolated from cells with RNeasy kit with Qiashredder cell disruption (Qiagen) and RNase-free DNase Set (Qiagen) on-column DNA digestion. DNA purity and concentration were assessed using an Agilent 2100 Bioanalyzer RNA Analysis chip (Eukaryote Total RNA Pico Series II) (Agilent Technologies). Samples from Exp. 1 underwent transcriptomic analysis using array-based gene expression analysis and samples from Exp. 2 using RNAseq.

For array-based gene expression analysis: Analysis of gene expression was performed using the MouseWG-6 v2.0 array (Illumina), following quality testing of mRNA using an Agilent 2100 Bioanalyzer. Data were analyzed in GeneSpring GX using two-way ANOVA and the multiple testing correction method of Benjamini–Hochberg. A fold change cutoff of ≥ ±2 was used to generate downstream data sets.

For RNAseq: Preparation of RNA library (mRNAlibrary preparation (poly A enrichment) and transcriptome sequencing (NovaSeq PE150 (6 G raw data per sample) was conducted by Novogene Co., LTD, using the HiSeq 2500 platform (Illumina) with a pair-end 150-nucleotide read length as described^4^. Raw sequencing data was mapped to the genome, raw and normalized counts (TPM) were determined, and differential gene expression was calculated using the CLC Genomic Workbench (Qiagen). Genes with adjusted p-value (q) < 0.05 and log(FoldChange) > 0.301 were considered as differentially expressed. Heatmaps and PC analyses/plots were generated using *ClustVis* ^5^.

Differentially expressed genes were used as input for Ingenuity IPA (Qiagen) to identify significantly regulated networks and pathways. Microbiome analysis: A 1 cm section of liquid nitrogen snap-frozen terminal ileum was used to quantify the mouse intestinal microbiome because of the advantages in quantifying the adherent mucosal microbiota^6^. We used ZymoBIOMICS whole organism and DNA microbial community standards (Zymo Research) as well as sterile-filtered PBS to create pairs of positive and negative controls appropriate for each step of the sample collection and isolation procedure. Microbial DNA was extracted and sequenced using the MiSeq platform (Illumina) at the UAB Microbiome Core using 16S V3-4 (bacterial) using primers 515F-806R.

#### Bioinformatics

We used QIIME 2.0 to merge, cluster, and quality control 16S rRNA gene amplicon sequences. Operational taxonomic units (OTUs) were called based on the SILVA database using closed reference-based OTU clustering with PyNAST and taxonomy via UCLUST. OTU tables were exported into R where they were normalized, transformed and analyzed with the following packages: *vegan, tidyverse, phyloseq, DESeq2, Maaslin2, SpiecEasi*, and *caret*. Two samples were excluded for insufficient read depth (below 1000 OTUs). OTUs present in <10% of samples were removed, and the remaining OTUs were transformed by total sum scaling. Alpha diversity using the raw OTU data was quantified with the Shannon and Chao1 Indices and significance testing was performed using Mann-Whitney U and a mixed linear model. We described beta diversity with principal components analysis of Euclidean distances and performed significance testing using PERMANOVA and PERMDISP. Differential abundance analysis was performed with DESeq2 and MaAsLin2. Feature selection was performed with Random Forest, utilizing the ranger package within caret.

#### Statistical analysis

Statistical analyses were performed in R, CLC Genomic Workbench or GraphPad Prism. In general, q < 0.05 after adjustment for multiple comparisons was considered statistically significant.

## Supplemental Information

**Figure S1.**
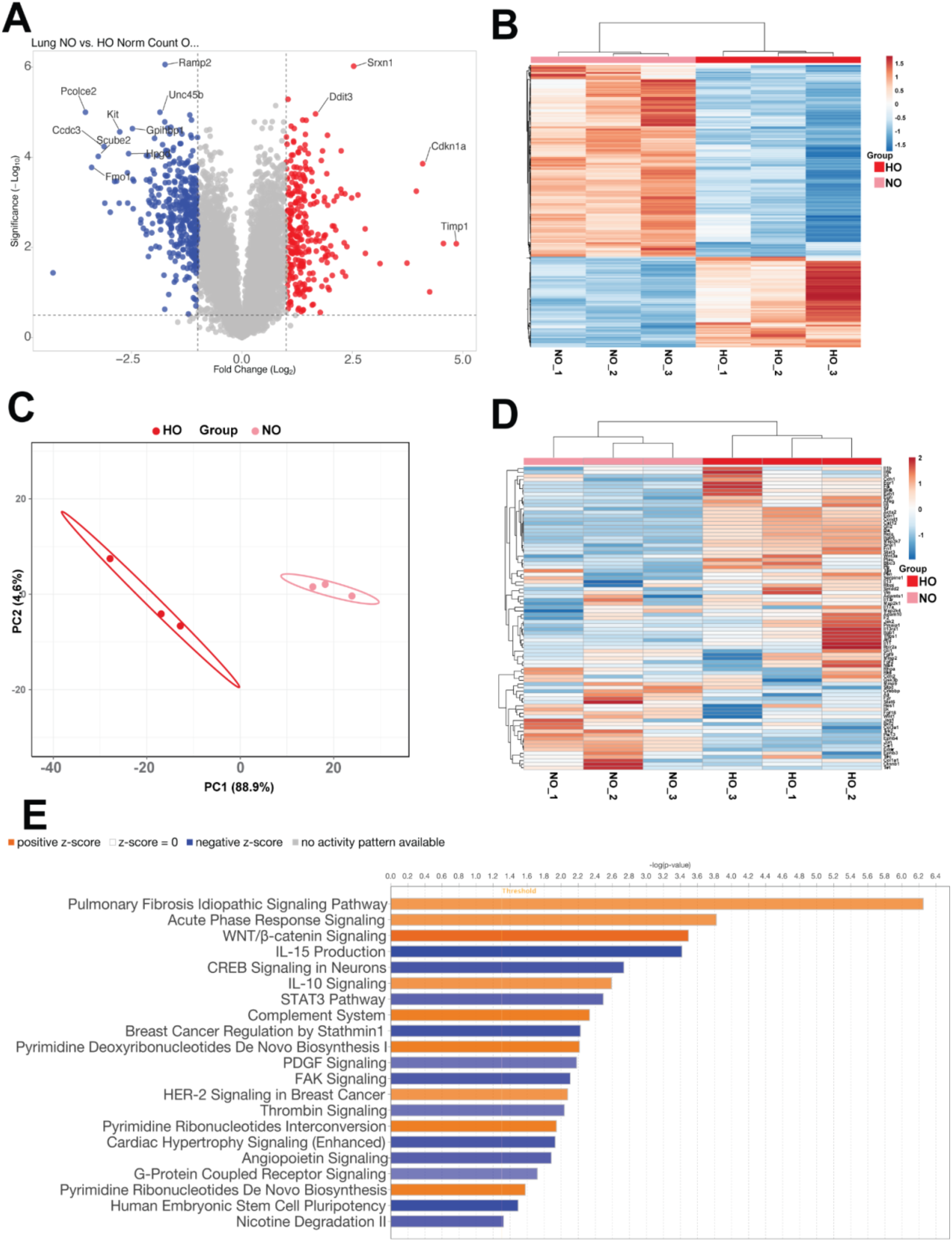
Hyperoxia exposure alters the mouse lung transcriptome. A) Volcano plot of lung gene expression array showing gene expression altered by hyperoxia exposure. NO, normoxia (pink). HO, hyperoxia (red). B) Heatmap showing differential gene expression in normoxia versus hyperoxia-exposed mice. C) Principal components analysis showing differential clustering of mice exposed to normoxia versus hyperoxia. PC, principal component. D) Heatmap showing differential expression of pulmonary fibrosis-related genes. E) Differentially expressed pathways in normoxia versus hyperoxia exposure, showing marked increases in fibrosis, acute phase response and WNT/β-catenin-related pathways.

**Figure S2.**
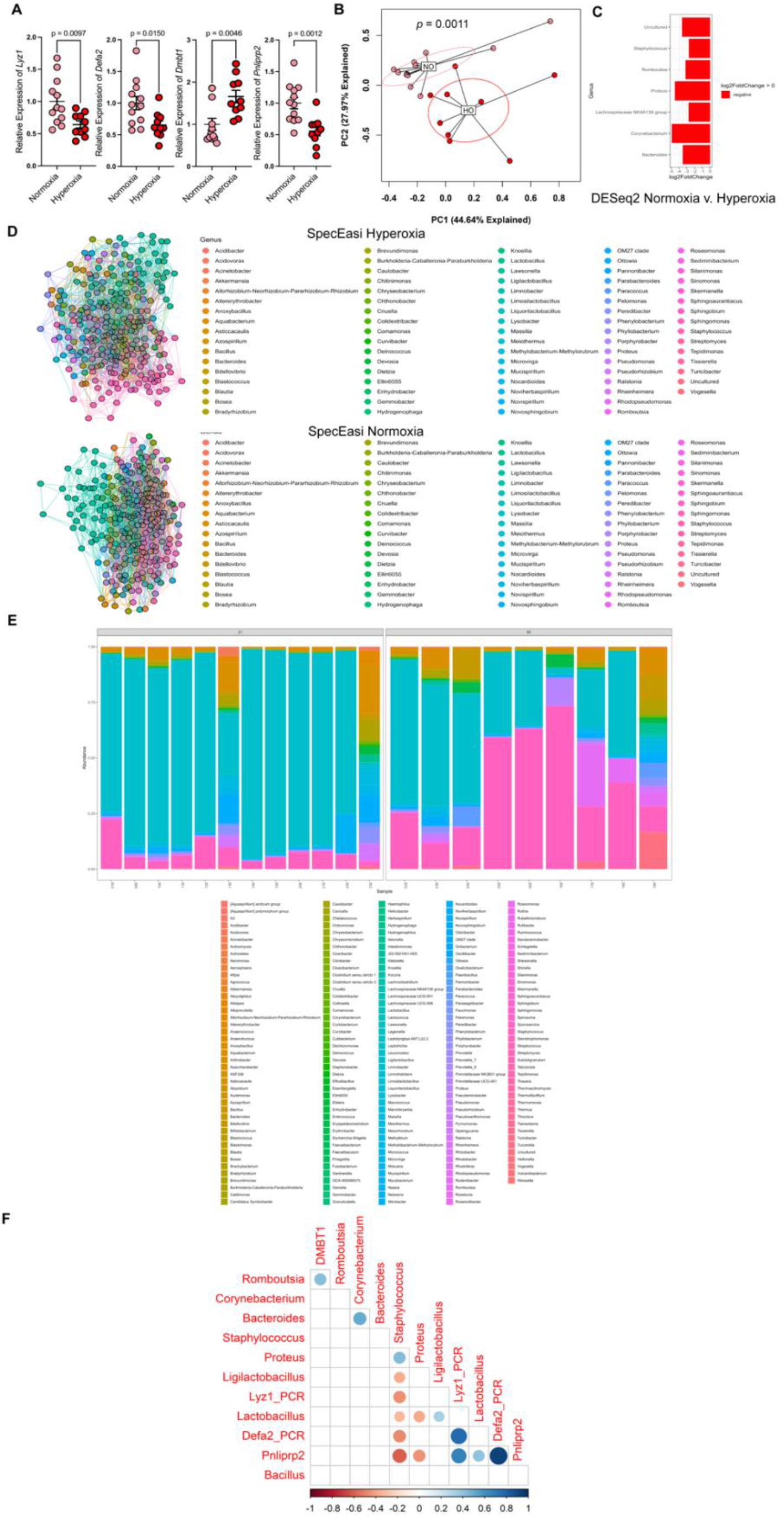
Hyperoxia alters the ileal microbiome. A) Quantitative PCR for a selection of differentially expressed genes showing alterations in expression of antimicrobial peptides and Paneth cell markers. B) Principal components analysis of Hellinger transformed Euclidian distances, p = 0.0011 pseudo-F = 5.2727 PERMANVOA, p = 1 PERMDISP, showing different global community composition between normoxia- and hyperoxia-exposed neonatal mice. The loading plot for this PCA is shown in Figure 1J. C) Binomial regression using DESeq2 showing differences at the genera level in normoxia- or hyperoxia-exposed animals. D) SpiecEasi network analysis of the microbiota of hyperoxia and normoxia exposed mice. E) Relative abundance at the genera level. F) Spearman correlation analysis showing association between antimicrobial peptide expression by qPCR and differentially abundant genera.

**Figure S3.**
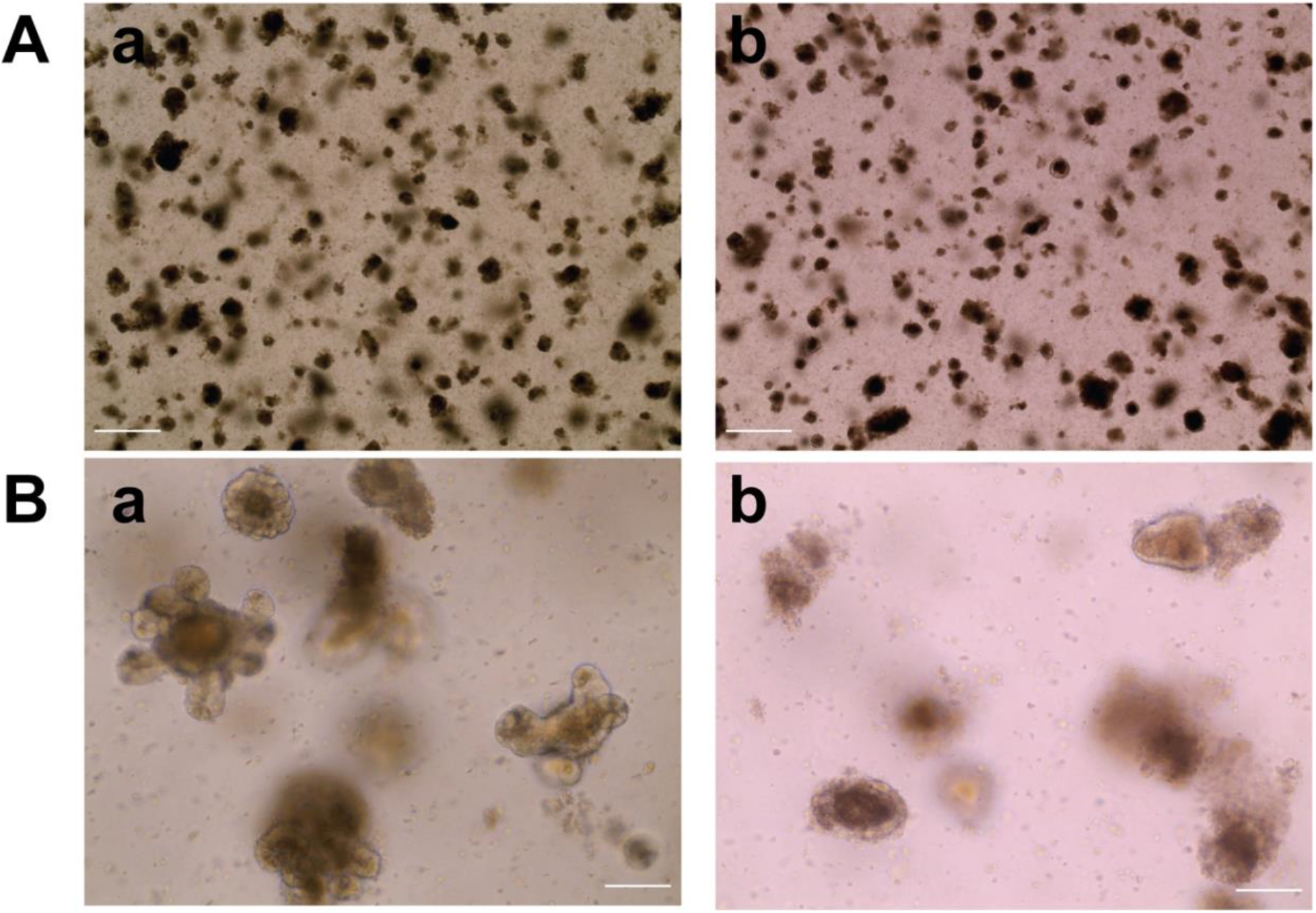
Hyperoxia exposure alters the morphology of intestinal organoids. A) Visual light microscopy at 4x of intestinal organoids after exposure to (a) normoxia or (b) hyperoxia for 24 h. Scale bar represents 1000 µm. B) Visual light microscopy and 20x magnification showing less budding and differences in epithelial integrity in (b) hyperoxia exposed organoids. Scale bar represents 200 µm.

**Figure S4.**
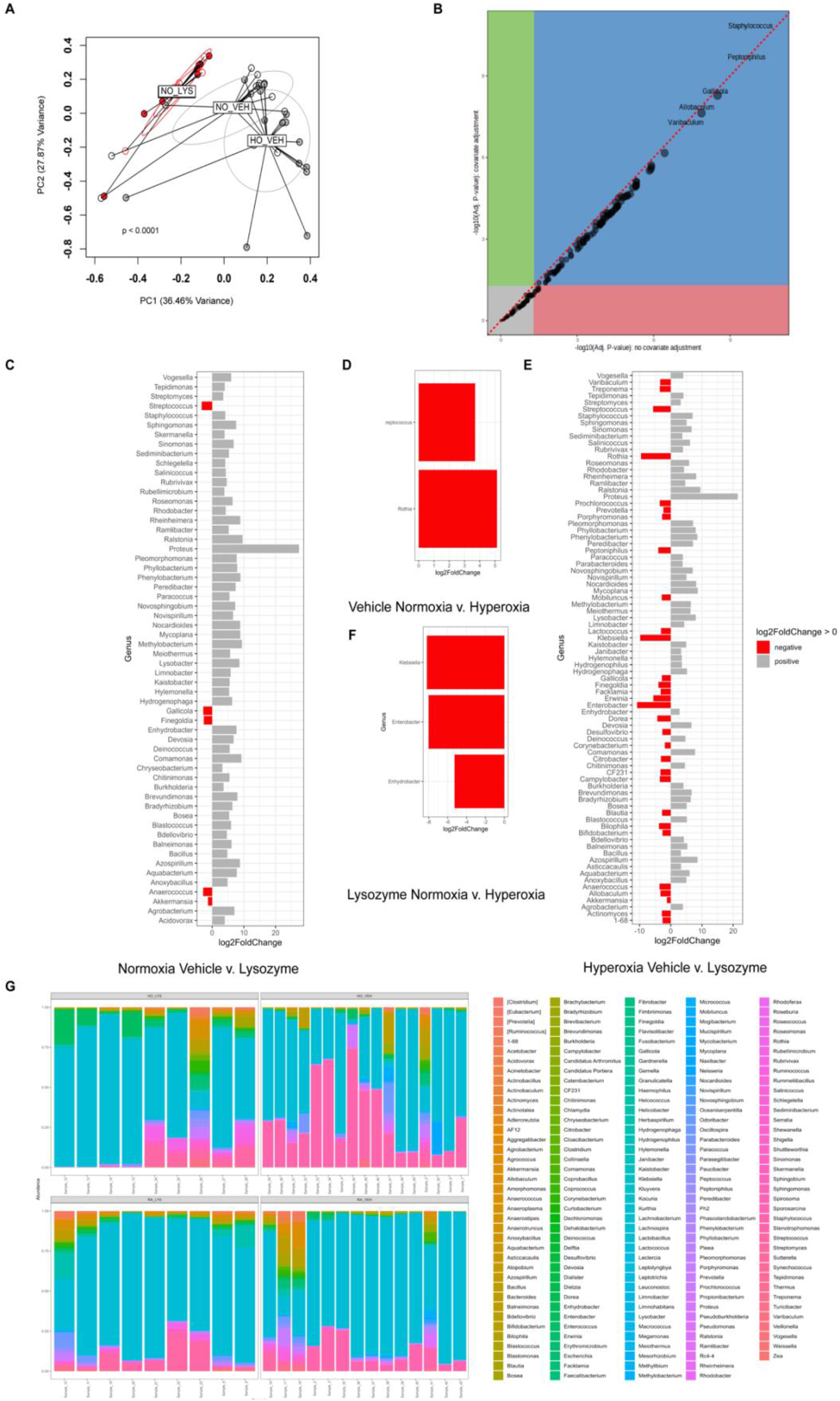
Lysozyme and hyperoxia alter the ileal microbiome. A) Principal components analysis of Hellinger transformed Euclidean distances showing significant differences in global community composition, p < 0.0001 PERMANOVA, p = 0.6524 PERMDISP. That is driven by differences between hyperoxia- and hyperoxia and lysozyme-exposed mice. (p = 0.0027, pairwise PERMANOVA hyperoxia versus hyperoxia and lysozyme). B) MaAsLin 2 (Microbiome Multivariable Associations with Linear Models) showing differences in the intestinal microbiota. C) Binomial regression showing differences at the genera level in normoxia between vehicle- and lysozyme exposed animals. D) Binomial regression showing differences between normoxia and hyperoxia vehicle-exposed control animals. E) Binomial regression showing differences between normoxia and hyperoxia lysozyme-exposed animals. F) Binomial regression showing differences at the genera level in hyperoxia between vehicle- and lysozyme exposed animals. G) Relative abundance at the genera level.

**Table S1.**
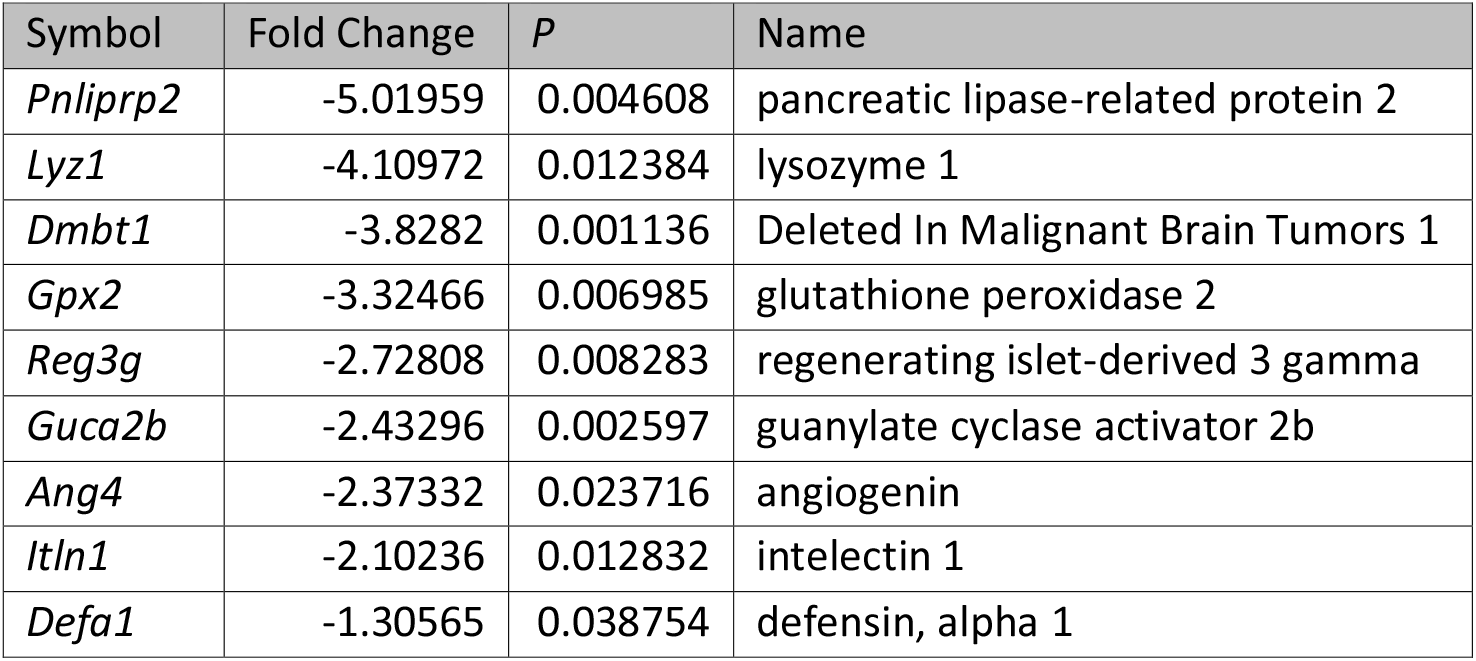
Paneth cell markers significantly changed in the murine ileum in response to hyperoxia.

**Table S2.** Paneth cell markers significantly changed in the murine intestinal organoids in response to hyperoxia.

| Symbol | Fold Change | P | Name |
| --- | --- | --- | --- |
| *Mptx2* | -6.10435 | 2.47E-06 | mucosal pentraxin 2 |
| *Dmbt1* | -5.79307 | 0.027013 | deleted in malignant brain tumors 1 |
| *Defa26* | -3.80301 | 1.77E-06 | defensin, alpha, 26 |
| *Kcnn4* | -2.89266 | 3.81E-05 | potassium intermediate/small conductance calcium-activated channel, subfamily N, member 4 |
| *Defa23* | -2.60226 | 0.000873 | defensin, alpha, 23 |
| *Ang4* | -2.38428 | 0.003666 | angiogenin, ribonuclease A family, member 4 |
| *Defa20* | -2.34975 | 0.013918 | defensin, alpha, 20 |
| *Defa22* | -2.2797 | 0.023124 | defensin, alpha, 22 |
| *Defa21* | -2.26413 | 0.027105 | defensin, alpha, 21 |
| *Defa5* | -2.20591 | 0.029043 | defensin, alpha, 5 |
| *Wnt3* | -1.70433 | 0.049799 | wingless-type MMTV integration site family, member 3 |
| *Spink4* | -1.65614 | 0.043295 | serine peptidase inhibitor, Kazal type 4 |
| *Pnliprp2* | -1.63384 | 0.000914 | pancreatic lipase-related protein 2 |
| *Dll4* | -1.54349 | 0.040336 | delta like canonical Notch ligand 4 |
| *Rap1a* | 1.351647 | 0.054885 | RAS-related protein 1a |
| *Tm4sf20* | 1.902737 | 0.001682 | transmembrane 4 L six family member 20 |
| *Clca1* | 2.660606 | 0.000193 | chloride channel accessory 1 |
| *Guca2b* | 3.273845 | 1.09E-06 | guanylate cyclase activator 2b (retina) |
| *Guca2a* | 3.743392 | 6.98E-09 | guanylate cyclase activator 2a (guanylin) |

**Table S3.** Lung inflammation-related genes as compared to normoxia vehicle exposed mice.

| Symbol | HO VEH vs. NO VEH |  | HO LYZ vs NO VEH |  | Database object name |
| --- | --- | --- | --- | --- | --- |
|  | Fold Change | q | Fold Change | q |  |
| *Ahsg* | 626.6391 | 0.005346 | 21.07616 | 0.269682 | alpha-2-HS-glycoprotein |
| *Csrp3* | 14.15701 | 0.000613 | -1.133 | 0.927459 | cysteine and glycine-rich protein 3 |
| *Mylk3* | 6.296584 | 0.000862 | -1.28659 | 0.76882 | myosin light chain kinase 3 |
| *Il1r2* | 3.475312 | 0.025753 | 1.949132 | 0.294827 | interleukin 1 receptor, type II |
| *Ptger3* | 2.537922 | 0.000685 | 1.6335 | 0.116541 | prostaglandin E receptor 3 (subtype EP3) |
| *Pbk* | 2.51287 | 0.017812 | 1.556554 | 0.330692 | PDZ binding kinase |
| *Ccl6* | 2.087173 | 4.42E-06 | 1.087488 | 0.740466 | chemokine (C-C motif) ligand 6 |
| *Clec7a* | 1.904633 | 4.27E-06 | -1.19229 | 0.341095 | C-type lectin domain family 7, member a |
| *Gpx1* | 1.797218 | 2.05E-06 | 1.19287 | 0.262587 | glutathione peroxidase 1 |
| *Ccr2* | 1.754312 | 0.026097 | 1.488087 | 0.138412 | chemokine (C-C motif) receptor 2 |
| *Irf5* | 1.608403 | 0.014589 | 1.329064 | 0.186512 | interferon regulatory factor 5 |
| *Chid1* | 1.593086 | 0.006607 | 1.357239 | 0.100582 | chitinase domain containing 1 |
| *Ndufs4* | 1.580271 | 0.000592 | 1.035841 | 0.872654 | NADH:ubiquinone oxidoreductase core subunit S4 |
| *Ctla2a* | 1.517331 | 0.00679 | 1.172581 | 0.402791 | cytotoxic T lymphocyte-associated protein 2 alpha |
| *Stk39* | 1.513097 | 0.043299 | 1.442597 | 0.074579 | serine/threonine kinase 39 |
| *Bst1* | 1.51008 | 0.035102 | 1.110652 | 0.693385 | bone marrow stromal cell antigen 1 |
| *Dicer1* | -1.58638 | 0.000146 | -1.1519 | 0.361353 | dicer 1, ribonuclease type III |
| *F2r* | -1.67139 | 3.58E-05 | -1.13661 | 0.444461 | coagulation factor II (thrombin) receptor |
| *Ackr2* | -1.69257 | 0.00167 | -1.08879 | 0.730939 | atypical chemokine receptor 2 |
| *Kdm6b* | -1.6956 | 2.7E-05 | -1.29584 | 0.071318 | KDM1 lysine (K)-specific demethylase 6B |
| *Akna* | -1.73878 | 0.001889 | -1.44747 | 0.052764 | AT-hook transcription factor |
| *Cybb* | -1.75339 | 0.001212 | -1.36063 | 0.113292 | cytochrome b-245, beta polypeptide |
| *Cdh5* | -1.78237 | 2.99E-05 | -1.12635 | 0.542984 | cadherin 5 |
| *Adora2a* | -2.30738 | 0.000152 | -1.4867 | 0.113343 | adenosine A2a receptor |
| *C1qtnf3* | -2.98067 | 0.006724 | -1.60992 | 0.30865 | C1q and tumor necrosis factor related protein 3 |

**Table S4.** Genes-related to endothelial cell regulation.

|  | HO VEH vs. RA VEH |  | HO LYZ vs RA VEH |  |  |
| --- | --- | --- | --- | --- | --- |
| Symbol | FC | q | FC | q | Database object name |
| *Mt3* | 171.49 | 0.039911 | 79.74 | 0.085312 | metallothionein 3 |
| *Gata4* | 3.57 | 0.001527 | 1.45 | 0.48924 | GATA binding protein 4 |
| *Igf2* | 2.56 | 8.07E-05 | -1.22 | 0.560584 | insulin-like growth factor 2 |
| *Aldh1a2* | 2.41 | 2.33E-05 | 1.19 | 0.560409 | aldehyde dehydrogenase family 1, subfamily A2 |
| *Smoc2* | 2.10 | 1.53E-05 | 1.08 | 0.785722 | SPARC related modular calcium binding 2 |
| *Gpx1* | 1.80 | 2.05E-06 | 1.19 | 0.262587 | glutathione peroxidase 1 |
| *Ccr2* | 1.75 | 0.026097 | 1.49 | 0.138412 | chemokine (C-C motif) receptor 2 |
| *Itgb1bp1* | 1.69 | 0.000516 | 1.25 | 0.216524 | integrin beta 1 binding protein 1 |
| *Nr4a1* | 1.67 | 0.034821 | -1.17 | 0.62015 | nuclear receptor subfamily 4, group A, member 1 |
| *Mydgf* | 1.60 | 0.000569 | 1.32 | 0.060535 | myeloid derived growth factor |
| *Fgf18* | 1.50 | 0.013835 | 1.03 | 0.916387 | fibroblast growth factor 18 |
| *Stard13* | -1.50 | 0.025186 | -1.04 | 0.9001 | StAR-related lipid transfer (START) domain containing 13 |
| *Jup* | -1.50 | 0.003187 | -1.18 | 0.324177 | junction plakoglobin |
| *Cxcl12* | -1.50 | 0.01231 | 1.01 | 0.985026 | chemokine (C-X-C motif) ligand 12 |
| *Paxip1* | -1.55 | 0.005012 | -1.24 | 0.22634 | PAX interacting (with transcription-activation domain) protein 1 |
| *Crkl* | -1.56 | 0.003138 | -1.03 | 0.909193 | v-crk avian sarcoma virus CT10 oncogene homolog-like |
| *Dicer1* | -1.59 | 0.000146 | -1.15 | 0.361353 | dicer 1, ribonuclease type III |
| *Mecp2* | -1.59 | 0.000473 | -1.08 | 0.682247 | methyl CpG binding protein 2 |
| *Jag1* | -1.59 | 0.002791 | 1.14 | 0.53448 | jagged 1 |
| *Flt1* | -1.59 | 0.008216 | 1.03 | 0.929469 | FMS-like tyrosine kinase 1 |
| *Ece1* | -1.64 | 1.62E-05 | -1.19 | 0.207628 | endothelin converting enzyme 1 |
| *Fzd4* | -1.67 | 0.034973 | 1.25 | 0.431742 | frizzled class receptor 4 |
| *Atoh8* | -1.68 | 0.001662 | -1.39 | 0.06783 | atonal bHLH transcription factor 8 |
| *Kdm6b* | -1.70 | 2.7E-05 | -1.30 | 0.071318 | KDM1 lysine (K)-specific demethylase 6B |
| *Ptprm* | -1.71 | 2.19E-05 | -1.06 | 0.778646 | protein tyrosine phosphatase, receptor type, M |
| *Sp1* | -1.72 | 4.56E-05 | -1.20 | 0.26853 | trans-acting transcription factor 1 |
| *Cdh5* | -1.78 | 2.99E-05 | -1.13 | 0.542984 | cadherin 5 |
| *Pecam1* | -1.79 | 2.22E-05 | -1.23 | 0.220446 | platelet/endothelial cell adhesion molecule 1 |
| *Pde4d* | -1.81 | 0.000142 | -1.33 | 0.104711 | phosphodiesterase 4D, cAMP specific |
| *Flt4* | -1.87 | 1.53E-06 | -1.18 | 0.332498 | FMS-like tyrosine kinase 4 |
| *Il6ra* | -2.55 | 4E-05 | -1.56 | 0.084073 | interleukin 6 receptor, alpha |
| *Lgals12* | -5.58 | 0.043358 | 1.26 | 0.832064 | lectin, galactose binding, soluble 12 |
| *Fgf2* | -6.15 | 0.047026 | 2.32 | 0.342984 | fibroblast growth factor 2 |

**Table S5.** Genes with opposite regulation associated with lysozyme supplementation.

|  | HO_VEH<br>vs.<br>RA_VEH<br>FC | HO_VEH<br>vs.<br>RA_VEH q | HO_LYZ<br>vs.<br>RA_VEH<br>FC | HO_LYZ<br>vs.<br>RA_VEH<br>q | HO_LYZ<br>vs.<br>HO_VEH<br>FC | HO_LYZ<br>vs.<br>HO_VEH<br>q | Database object name |
| --- | --- | --- | --- | --- | --- | --- | --- |
| *Sln* | 13.69 | 0.000426 | -11.55 | 0.003672 | -158.08 | 2.94E-10 | sarcolipin |
| *Myl7* | 6.38 | 0.001178 | -4.37 | 0.013565 | -27.89 | 4.11E-09 | myosin, light polypeptide 7, regulatory |
| *Hba-a2* | 4.21 | 0.000184 | -2.45 | 0.032445 | -10.30 | 1.53E-09 | hemoglobin alpha, adult chain 2 |
| *Hbb-bt* | 4.16 | 0.000119 | -2.83 | 0.007483 | -11.79 | 2.44E-11 | hemoglobin, beta adult t chain |
| *Hba-a1* | 3.87 | 0.000579 | -2.65 | 0.019222 | -10.22 | 2.74E-09 | hemoglobin alpha, adult chain 1 |
| *Snca* | 3.63 | 0.000852 | -2.38 | 0.038694 | -8.65 | 2.51E-08 | synuclein, alpha |
| *Alas2* | 3.61 | 0.000565 | -3.21 | 0.001881 | -11.59 | 2.27E-11 | aminolevulinic acid synthase 2, erythroid |
| *Hbb-bs* | 2.97 | 0.016819 | -4.08 | 0.000942 | -12.09 | 4.03E-09 |  |
| *Slc4a1* | 2.47 | 0.008169 | -2.32 | 0.015472 | -5.74 | 1.53E-07 | solute carrier family 4 (anion exchanger), member 1 |
| *Cpe* | 1.74 | 0.000235 | -1.75 | 0.000201 | -3.03 | 5.23E-14 | carboxypeptidase E |
| *Igfbp5* | 1.51 | 0.041052 | -1.82 | 0.001169 | -2.75 | 4.24E-08 | insulin-like growth factor binding protein 5 |
| *mt-Rnr2* | 1.51 | 0.016924 | -1.60 | 0.004611 | -2.41 | 5.45E-08 | mitochondrially encoded 16S rRNA |
| *S100a1* | 1.39 | 0.036962 | -1.41 | 0.023687 | -1.97 | 7.7E-06 | S100 calcium binding protein A1 |
| *Glg1* | -1.53 | 0.034846 | 1.86 | 0.000484 | 2.84 | 6.73E-09 | golgi apparatus protein 1 |
| *Abhd2* | -1.99 | 0.003723 | 1.79 | 0.014075 | 3.56 | 3.21E-08 | abhydrolase domain containing 2 |

